# Widespread circulation of West Nile and Usutu viruses in sedentary and migratory avifauna: A Two-Year Study (2024–2025) of active surveillance in South of France

**DOI:** 10.64898/2026.05.21.726855

**Authors:** Rachel Beaubaton, Justine Revel, Laetitia Pigeyre, Alexandre Lepeule, Julien Joly, Christophe de Franceschi, Anne Charmantier, Benjamin Vollot, Yannick Simonin

## Abstract

West Nile virus (WNV) and Usutu virus (USUV) are neurotropic Orthoflaviviruses sharing a similar enzootic transmission cycle primarily involving Culex pipiens mosquitoes as vectors and birds as amplifying hosts. First identified in Africa, both viruses established endemicity across Europe over the past two decades, most likely introduced and spread by migratory bird species along Mediterranean flyways. In avian species, infection outcomes range from subclinical to fatal neuroinvasive disease, varying by viral strain, host immunity, and species susceptibility. Southern France emerges as a key hotspot for the circulation of these viruses, supported by diverse avian habitats conducive to year-round viral maintenance. This study investigated the prevalence of WNV and USUV in more than 2500 sedentary and migratory wild birds from these regions during 2024–2025 using molecular surveillance. Samples were collected using mist net and bird boxes, across multiple passerine and non-passerine taxa, spanning wetlands, urban fringes, and agricultural zones. Our analyses revealed widespread viral circulation across diverse species, mainly among passerines such as great tits, house sparrows, and barn swallows with USUV detected at higher rates than WNV in both study years. Overall prevalence was markedly higher in 2024 than in 2025, potentially reflecting climatic or ecological drivers. Migratory individuals likely seed viral introductions during seasonal passages, whereas resident populations sustain local enzootic cycles, facilitating overwintering persistence. These results highlight the pivotal role of mixed avifauna in arbovirus dynamics within Mediterranean Europe and emphasize the necessity for integrated, year-round surveillance targeting high-risk species and habitats. Enhanced monitoring will aid in predicting spillover risks and informing vector control strategies to mitigate zoonotic threats.

## Introduction

West Nile virus (WNV) and Usutu virus (USUV), closely related members of the Japanese encephalitis virus serocomplex, are arboviruses of the genus *Orthoflavivirus* from the *Flaviviridae* family. They were first identified and isolated in Africa, WNV in 1937 in Uganda in the West Nile district (1) and USUV in 1959 in South Africa near the Usutu River (2). They share similar biological traits, transmission cycle and epidemiology (3). *Culex pipiens* mosquitoes, which are primarily ornithophilic, can transmit WNV and USUV from birds, considered the main amplifying hosts, to various mammalian species, including humans, horses, dogs, bats, wild boars, and several ruminants such as deer, cattle, and sheep, which are presumed to be dead-end hosts (4,5). Most human infections with WNV and USUV are asymptomatic, but when symptoms occur, they are usually mild, including fever, headache, rash, or muscle and joint pain. Severe neurological complications, including encephalitis or meningitis, may occasionally occur. Current evidence suggests that WNV is more virulent in humans than USUV whereas the opposite pattern is observed in birds (1,5–8). Approximately 30 neuroinvasive USUV cases of varying severity in humans have been reported in Europe since 2016, while WNV occasions each year several hundred human cases and dozens of deaths across Europe (4,6). USUV is classified into eight distinct genetic lineages, grouped into African (Africa 1/2/3) and European (Europe 1/2/3/4/5) clusters, with virulence potentially varying between lineages (9–11). Among the nine known WNV lineages, infections are primarily caused by lineages 1 and 2, with lineage 2 notably contributing to the recent rise in severe neuroinvasive human cases in Europe (12,13).

The epidemiology of WNV and USUV has changed considerably over the past two decades in Europe, with repeated epizootic events contributing to their progressive endemization. USUV was first associated with significant avian mortality in Europe in 1996, affecting blackbird (*Turdus merula*) populations in Italy (6). A notable amplification event occurred in 2018, when outbreaks of both viruses affected several European countries (4,14–16). USUV was associated with high mortality in multiple bird species, particularly blackbirds, magpies (*Pica pica*) and Strigiformes. During the same season, WNV activity also increased in avian hosts, with initial detections in resident wild and captive birds in Germany and widespread evidence of WNV infection across multiple bird species and carcasses (4,16–20). Since then, both viruses have become widely established across Central and Western Europe, causing recurrent avian outbreaks and demonstrating a progressive northward expansion of their distribution range (21–29). The endemization of WNV and USUV in Europe is thought to be influenced by repeated introductions mediated by long-distance migratory birds along major Afro-Palearctic flyways, followed by local amplification within resident bird–mosquito transmission cycles (30). This applies to tits (*Paridae*), sparrows (*Passeridae*) and pigeons (*Columbidae*), which are resident passerines widely distributed across the continent and identified in the literature as being susceptible to both WNV and USUV (7,31). Migratory species proposed as potential contributors to viral dispersal into new European regions include the common kestrel (*Falco tinnunculus*) and the lesser whitethroat (*Curruca curruca*), both of which migrate between Africa and Europe (4–6,9). The first recorded entry from Africa into Spain aligns with the East Atlantic migratory pathway, while its introduction to Central Europe appears to follow the Black Sea/Mediterranean route (6). Comparable introduction patterns have been described for WNV, with multiple lineage exchanges between Africa and Europe along these migratory corridors (32).

In France, WNV and USUV infections have mainly been reported in the south, especially in the city of Montpellier. The nearby Camargue Regional Nature Park and surrounding wetlands provide an optimal environment for arbovirus maintenance, with dense avian populations, proximity to major migratory routes, and high mosquito vector activity (33). Infections have been reported in humans (34), mosquitoes (33,35), birds (36,37), dogs, horses (33) and in several species of the Montpellier zoological park (25,37). Notably, the first human USUV infection in France was reported in 2016, in a patient from Montpellier presenting idiopathic facial paralysis (34). Serological screening in the same region revealed that 3% of blood donors had antibodies against USUV, suggesting local viral circulation (33). Moreover, the first human case of WNV in Europe was identified in the Camargue in 1962, and in 2024 12 human cases of WNV infection were reported in our study area (38).

WNV and USUV primarily circulate in wild birds and follow complex transmission cycles, making surveillance of both resident and migratory species essential to understand their dynamics. Accordingly, in 2024 and 2025, we implemented a monitoring program in the Montpellier-Camargue region to identify the bird species most susceptible to infection, with the goal of better assessing virus circulation and anticipating potential epizootics in humans and equines. More than 2500 avian samples, across 69 species, were sampled between May–December 2024 and March–November 2025 at different sites (based on avian density, wetlands proximity, and vector density). Cloacal swabs and droppings were analyzed using a WNV/USUV RT-qPCR TaqMan duplex assay. Our findings reveal a markedly high circulation of both viruses in avian hosts in 2024, followed by lower activity in 2025. This circulation occurred early in the season, emerging well before the detection of cases in humans and horses, and even preceding virus detection in mosquito vectors. Over the two-year period, USUV exhibited nearly twice the molecular prevalence of WNV across both resident and migratory bird species, with limited co-infections. Several species showed particularly high prevalence for both viruses, most notably great tits (*Parus major)*, house sparrows (*Passer domesticus)*, and barn swallows (*Hirundo rustica)*. Viral activity was detected across urban, peri-urban, and rural habitats, underscoring the broad ecological footprint of both pathogens. Together, these patterns highlight the central role of avifauna in the early amplification and spatial dynamics of WNV and USUV.

## Methods

### Survey area

The study was conducted in southern France, within the Mediterranean basin, and focused on a geographically continuous region encompassing two main areas: the Camargue wetland and the urban and peri-urban surroundings of Montpellier, located in the eastern part of the Occitanie region. This region is characterized by a warm-summer Mediterranean climate as defined by the Köppen–Geiger classification (39). The Camargue represents one of the largest natural wetland complexes in Western Europe and is situated between the two arms of the Rhône River and the Mediterranean coastline. This study area is mainly located in the Petite Camargue in the Occitanie region, corresponding to the western part of the Camargue. The flat and low landscape is shaped by an extensive network of cultures, canals, ditches, and wetlands that maintain high levels of humidity even in otherwise dry areas. These environmental conditions support both diverse avian communities and dense mosquito populations, creating a favorable ecological context for arbovirus circulation (40). Birds were sampled in lowland areas, where the WNV and USUV vector *Culex pipiens* is most abundant. In 2024, birds were sampled in seven different municipalities (Figure 1A): 43°39’45.8”N 3°39’52.8”E (Montarnaud, 34163); 43°49’25.1”N 3°47’43.5”E (Rouet, 34380); 43°35’36.8”N 4°11’57.2”E (Tour Carbonnière, Saint-Laurent-d’Aigouze, 30220); 43°42’34.8”N 4°12’12.7”E (Carrière, Aigues-Vives, 30670); 43°35’04.4”N 4°12’27.6”E (Aigues-Mortes, 30220); 43°36’20.7”N 4°20’13.9”E (Centre de découverte du Scamandre, Vauvert, 30600); 43°38’42.4”N 4°24’14.0”E (Espeyran, Saint-Gilles, 30800). In 2025, 16 different municipalities were used for sampling (Figure 1B): 43°39’45.8”N 3°39’52.8”E (Montarnaud, 34163); 43°49’28.6”N 3°47’39.2”E (Rouet, 34380); 43°35’36.8”N 4°11’57.2”E (Tour Carbonnière, Saint-Laurent-d’Aigouze, 30220); 43°42’34.8”N 4°12’12.7”E (Carrière, Aigues-Vives, 30670); 43°35’04.4”N 4°12’27.6”E (Aigues-Mortes, 30220); 43°36’20.7”N 4°20’13.9”E (Centre de découverte du Scamandre, Vauvert, 30600); 43°38’42.4”N 4°24’14.0”E (Espeyran, Saint-Gilles, 30800); 43°38’20.0”N 3°51’42.1”E (CEFE, Montpellier, 34090); 43°36’30.5”N 3°42’36.6”E (Murviel-lès-Montpellier, 34570); 43°18’43.1”N 3°31’48.2”E (Réserve Naturelle du Bagnas, Agde, 34300); 43°31’02.6”N 3°50’04.8”E (Villeneuve-lès-Maguelone, 34750); 43°33’04.7”N 4°06’54.7”E (Grau-du-Roi, 30240); 43°28’41.4”N 4°23’38.5”E (Saintes-Maries-de-la-Mer, 13460); 43°39’45.99”N 4°28’47.87”E (Arles, 13200); 43°42’08.0”N 4°14’18.2”E (Le Cailar, 30740); 43°43’23.6”N 4°13’53.7”E (Codognan, 30920). These sites were selected based on bird density, proximity to wetlands and/or the density of mosquito vectors. Sampling sites were classified a priori into three locality types according to their dominant surrounding land-use context. Urban sites corresponded to continuous built-up areas and included Montpellier. Peri-urban sites corresponded to transitional areas at the interface between built-up environments and agricultural or natural habitats and included Agde, Aigues-Mortes, Arles, Codognan, Le Cailar, Le Grau-du-Roi, Montarnaud and Murviel-lès-Montpellier. Rural sites corresponded to areas dominated by wetlands, agricultural land or natural habitats with low building density and included Aigues-Vives, Rouet, Saintes-Maries-de-la-Mer, Saint-Gilles, Saint-Laurent-d’Aigouze, Vauvert and Villeneuve-lès-Maguelone.

**Figure 1:**
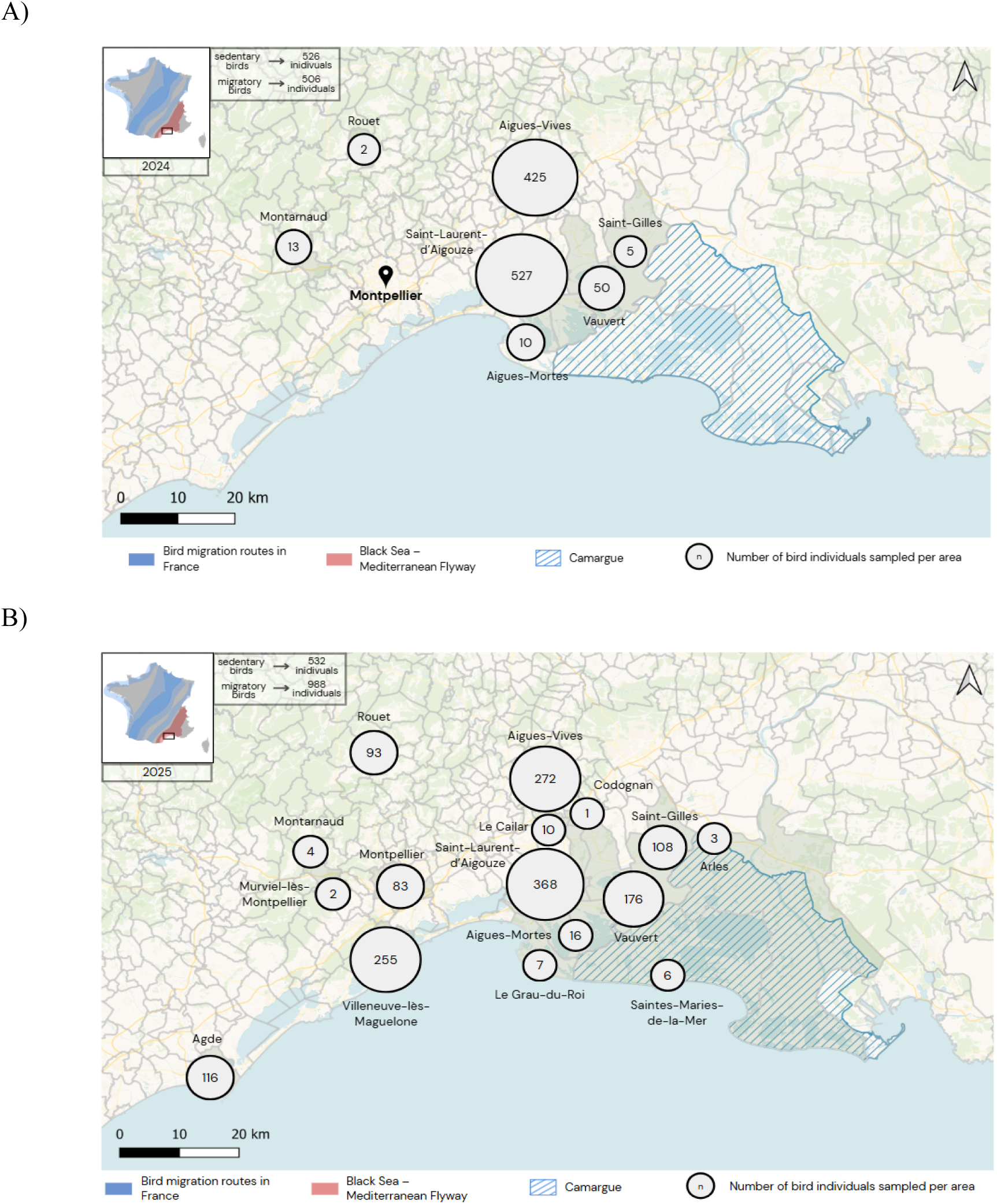
Maps illustrating the number of individual birds sampled at each capture site in 2024 (A) and 2025 (B).

### Spatiotemporal overview of avian sampling

In 2024, wild birds were captured from 15 May 2024 to 02 December 2024, with a total of 1110 samples from 1032 different individuals, including 54 bird species from 7 different orders. 506 individuals were migratory birds from 29 species, and 526 individuals were sedentary birds from 25 species. Of the 1110 samples, 1039 were cloacal swabs placed in RNA preservative medium, and 71 were droppings. In 2025, wild birds were captured from 23 February 2025 to 28 October 2025, with a total of 1773 samples from 1520 different individuals, including 60 bird species from 6 different orders. 988 individuals were migratory birds from 30 species, and 532 individuals were sedentary birds from 30 species. Of the 1773 samples, 1507 were cloacal swabs placed in RNA preservative medium, and 266 were droppings.

### Field capture protocol and biological sample collection procedures

Wild birds were captured using mist nets installed in habitats visited by the target species, particularly along flight corridors, forest edges, and near feeding areas. As the objective of this study was not to assess population trends, sampling effort was not standardized across sessions. For each session, the surface area covered, number of net-hours, and duration of net opening were recorded. Mist nets with an appropriate mesh size were selected according to the expected size of species present in the study area. Nets were mounted vertically between two poles and operated depending on weather conditions. They were checked at 30-minute intervals to minimize handling time, reduce stress, and limit the risk of injury to captured individuals. Captured birds were carefully removed from the nets and the species identified. Individuals were fitted with uniquely numbered metal leg bands provided by the Centre de Recherches sur la Biologie des Populations d’Oiseaux (CRBPO, French Museum of Natural History). All birds were released at their site of capture immediately after processing. Standard morphometric measurements were obtained from a subset of individuals, including body mass, wing length (flattened and straightened), tarsus length, and bill length. These checks are mandatory prior to sampling as they enable ornithologists to verify that the bird is in good health, but they are not used in our analyses. A visual clinical examination was performed on each individual. Cloacal swabs and fecal samples were collected following the standardized protocol described by Knutie et al. (41).

### RNA extraction, qRT-PCR and sequencing

RNA extraction was carried out from 140 µL of cloacal swabs and droppings using the QIAamp Viral RNA Mini Kit (Qiagen), according to the manufacturer’s instructions. For cloacal swabs, 140 µL of the RNA preservation medium in which the swab was immersed was directly processed. For droppings, samples were ground in 140 µL of PBS 1x prior to extraction. At the end of the procedure, 50 µL of RNA eluate was recovered for each sample before proceeding to a one-step duplex TaqMan qRT-PCR assay. After extraction, 5 μL of each RNA sample was tested with SuperScript™ IV (Invitrogen), WNV primers (Forward: CCTGTGTGAGCTGACAAACTTAGT and Reverse: GCGTTTTAGCATATTGACAGCC) with a probe (5’CY5-CCTGGTTTCTTAGACATCGAGATCT-3’BHQ2), and USUV primers (Forward: AAAAATGTACGCGGATGACACA and Reverse: TTTGGCCTCGTTGTCAAGATC) with a probe (5’FAM- CGGCTGGGACACCCGGATAACC-3’TAMRA). Cycling conditions were as follows: reverse transcription at 50°C for 15 minutes; inactivation of reverse transcriptase and activation of Taq polymerase at 95°C for 2 minutes; followed by 50 cycles of denaturation at 95°C for 15 seconds and elongation at 60°C for 1 minute. All reactions were performed on a LightCycler® 480 (Roche).

RT-qPCR–positive samples were subsequently subjected to a pan-orthoflavivirus PCR to generate larger amplicons suitable for downstream sequencing (42). RT-qPCR amplicons were purified using the NucleoSpin Gel and PCR Clean-up Kit and were subsequently submitted for sequencing. Sanger sequencing was performed by GENEWIZ (Azenta Life Sciences) using 20 µL of purified amplicons and the corresponding amplification primer.

### Phylogenetic analyses

For WNV thirty-three reference nucleotide sequences of the virus were aligned with four sequences from avian samples and three sequences from mosquitoes. The alignment was complemented by a sequence from Japanese encephalitis virus (JEV). For USUV fifty reference nucleotide sequences of the virus were aligned with two sequences from this study. The alignment was complemented by a sequence from WNV. The tool used for the alignment was MAFFT version 7.505 (_). The MAFFT output data were cleaned and trimmed using TrimAl version 1.5.rev1 (_) to optimize tree construction. A maximum likelihood phylogenetic tree was constructed with IQ-TREE version 3.0.1 (_), with automatic detection of the most appropriate model according to the Bayesian Information Criterion (BIC). Bootstrap percentages were obtained based on 1,000 replicates. The final tree was visualized and annotated with the online version iTOL version 7.5 (_) and rerooted with the JEV or WNV sequence.

### Ethical statement

All avian captures, handling, and sampling procedures were conducted in accordance with established ethical guidelines for wildlife research and were approved by the CRBPO, under the authority of the French National Museum of Natural History (Muséum national d’Histoire naturelle, MNHN; programme number 1331). Bird ringing permit No. 1820. For studies conducted by CEFE, authorization number F 3417211 was issued in February 2024. No animals were captured or sampled exclusively for the purpose of this study.

### Statistical analysis

Overall prevalence was calculated as the proportion of positive birds among the total number of birds tested and was reported with two-sided 95% confidence intervals (95% CI) using the Wilson method. Associations between virus positivity and categorical variables, including year and month of sampling, sex, age class, migratory status, Passeriformes status, feeding guild and locality type, were assessed using Pearson’s chi-square test when assumptions were met. When expected counts were below recommended thresholds, Fisher’s exact test was used instead. Differences in co-infection frequency between years were assessed using Fisher’s exact test. USUV and WNV positivity rates were compared using McNemar’s test.

To further assess factors associated with virus detection while accounting for potential confounding and species-level clustering, multivariable mixed-effects logistic regression models were fitted separately for USUV and WNV. Virus detection status was used as the binary outcome. For analyses, month of sampling was grouped into seasons as follows: winter, January-March; spring, April-June; summer, July-September; and autumn, October-December. Candidate explanatory variables included year, season, locality type, age, sex, migratory status, Passeriformes status and feeding guild. We first fitted univariate models with bird species included as a random effect. Final multivariate models were selected using log-likelihood ratio test criteria through nested model comparisons. Results were reported as odds ratios (OR) with 95% confidence intervals and p-values. The contribution of the species random effect was assessed using the intraclass correlation coefficient (ICC).

All statistical analyses were performed using R software (R Core Team, 2024) within the RStudio environment (version 4.4.2, 2024). A significance threshold of p < 0.05 was applied.

## Results

### Study areas and sampling strategies

Over the two-year study period, 2552 wild birds representing 69 species across eight taxonomic orders were sampled in urban, peri-urban, and rural areas (Supplementary Table 1; Figure 1).

In 2024, avian samples were collected from 1032 individual birds representing 54 species across seven orders. Among these species, 25 were sedentary species native to southern France, while the remaining 29 were migratory species (Supplementary Table 1). In 2025, 1520 individual birds were sampled, representing 60 species across six taxonomic orders, evenly distributed between sedentary (n = 30) and migratory species (n = 30). Compared with the 2024 campaign, 9 species were not sampled (*Actitis hypoleucos, Corvus corone, Ixobrychus minutus, Lanius collurio, Larus michahellis, Oriolus oriolus, Phylloscopus collybita tristis, Turdus philomelos, Upupa epops*), whereas 15 new species were sampled (*Acrocephalus paludicola, Caprimulgus europaeus, Certhia brachydactyla, Cisticola juncidis, Coccothraustes coccothraustes, Dryobates minor, Emberiza cirlus, Gallinago gallinago, Locustella naevia, Oenanthe oenanthe, Regulus ignicapilla, Saxicola rubetra, Strix aluco, Sylvia borin, Turdus merula*) (Supplementary Table 1).

In both years, Passeriformes constituted the largest proportion of species sampled (n = 56, 81.2%), with major contributions from barn swallow (*Hirundo rustica*) (n = 654, 25.6%), common reed warbler (*Acrocephalus scirpaceus*) (n = 223, 8.7%), great tit (*Parus major)* (n = 177, 6.9%) and house sparrow (*Passer domesticus)* (n = 149, 5.8%) (Figure 2). The remaining 18.8% of species were distributed across seven other orders, including Bucerotiformes, Caprimulgiformes, and Pelecaniformes as the most underrepresented, each being represented by a single species. Cloacal swabs and/or fecal samples were collected from each bird according to sampling opportunities. All samples were subsequently subjected to molecular analysis for the detection of USUV and WNV.

**Figure 2:**
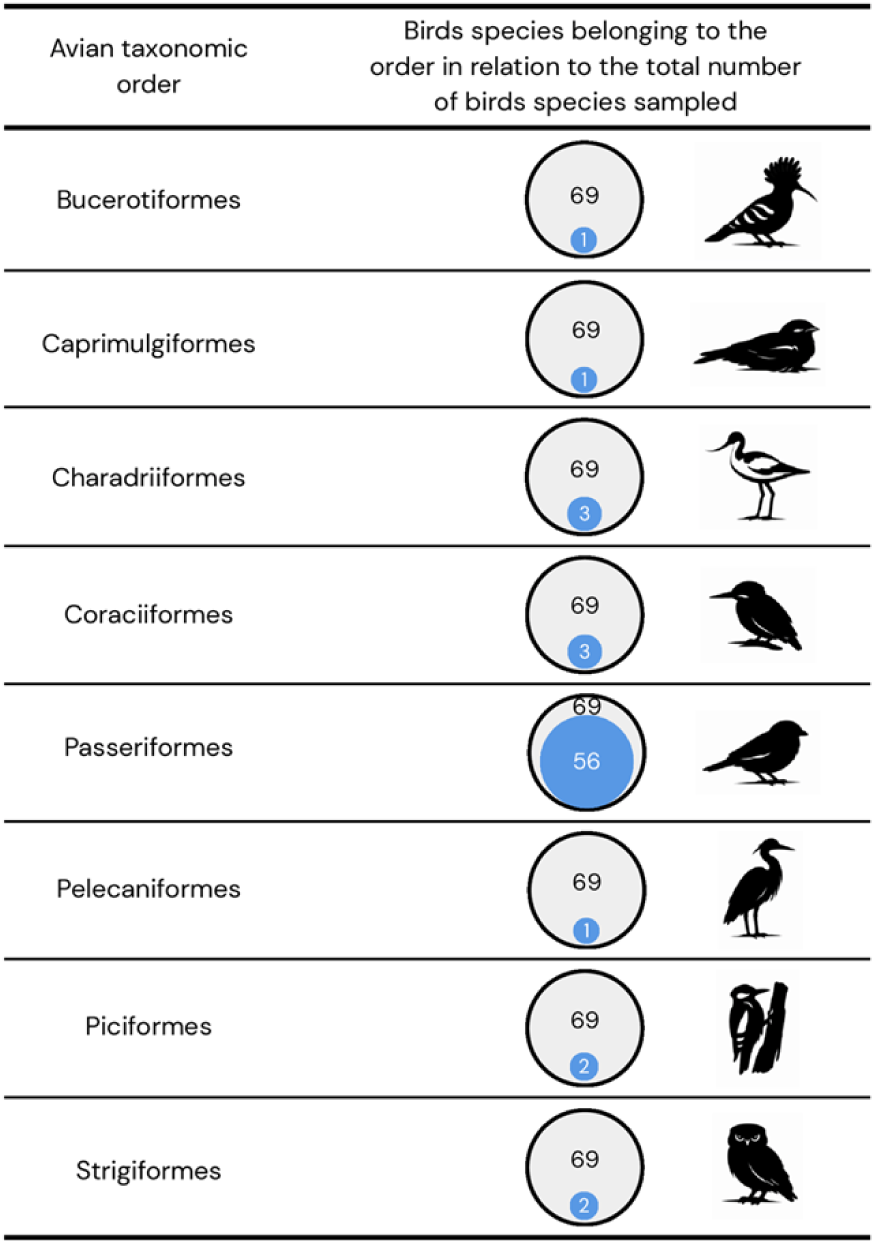
Proportion of avian taxonomic orders sampled in 2024 and 2025. Each row represents a bird taxonomic order. The grey circle indicates the total number of species sampled (n = 69, all orders combined). The blue circle represents the number of sampled species belonging to the focal order. Silhouettes on the right illustrate a representative species for each order.

Sedentary bird species are present in the study area throughout the year and are therefore potentially available for sampling across all seasons. In contrast, migratory species exhibit heterogeneous patterns of presence (Figure 3). Several migratory species are known to use the Camargue area as a breeding and resting site and typically remain in the region for several months between spring and autumn, resulting in prolonged local presence during the sampling period. For example, the common reed warbler (*Acrocephalus scirpaceus*), the European bee-eater (*Merops apiaster*), the barn swallow (*Hirundo rustica*) and the great reed warbler (*Acrocephalus arundinaceus*) arrive in spring to breed and remain throughout summer before departing in autumn. Other migratory species are mainly present during shorter stopover phases associated with spring or autumn migration, leading to more limited temporal windows of exposure. Examples include various species of warblers such as the sedge warbler (*Acrocephalus schoenobaenus)* or the willow warbler (*Phylloscopus trochilus*), which transit through the Camargue during their migration and are typically observed only during these seasonal windows. By contrast, truly sedentary species, such as the Eurasian blue tit (*Cyanistes caeruleus*), the great tit (*Parus major*), the moustached warbler (*Acrocephalus melanopogon*), or the house sparrow (*Passer domesticus*), are recorded throughout the year, consistent with their resident status in the local habitats. These differences in residency status and duration of presence align with the known migratory ecology of the species involved and may influence their exposure to locally circulating arboviruses, with prolonged residents having a longer potential contact period with virus vectors compared to short-distance or passage migrants.

**Figure 3:**
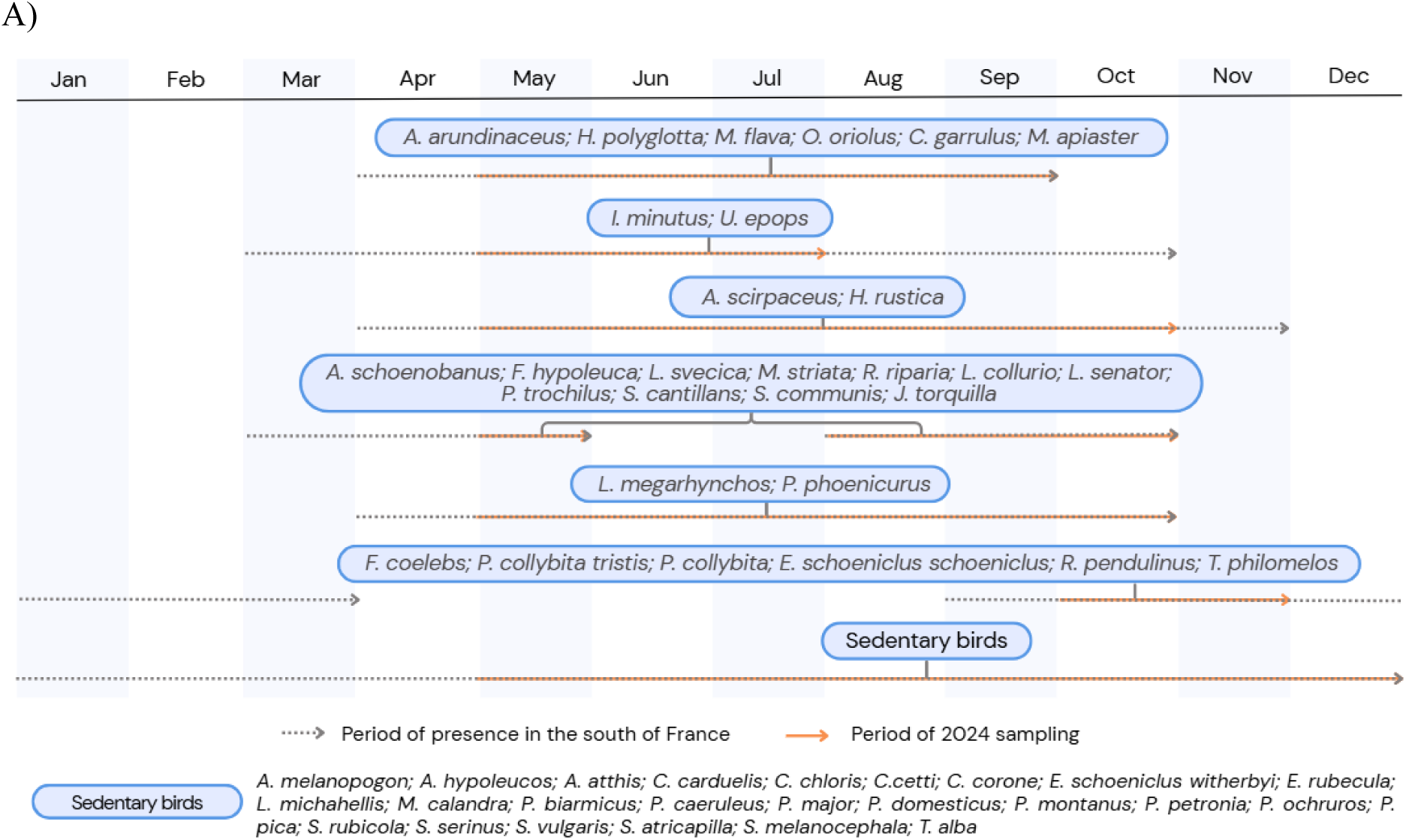
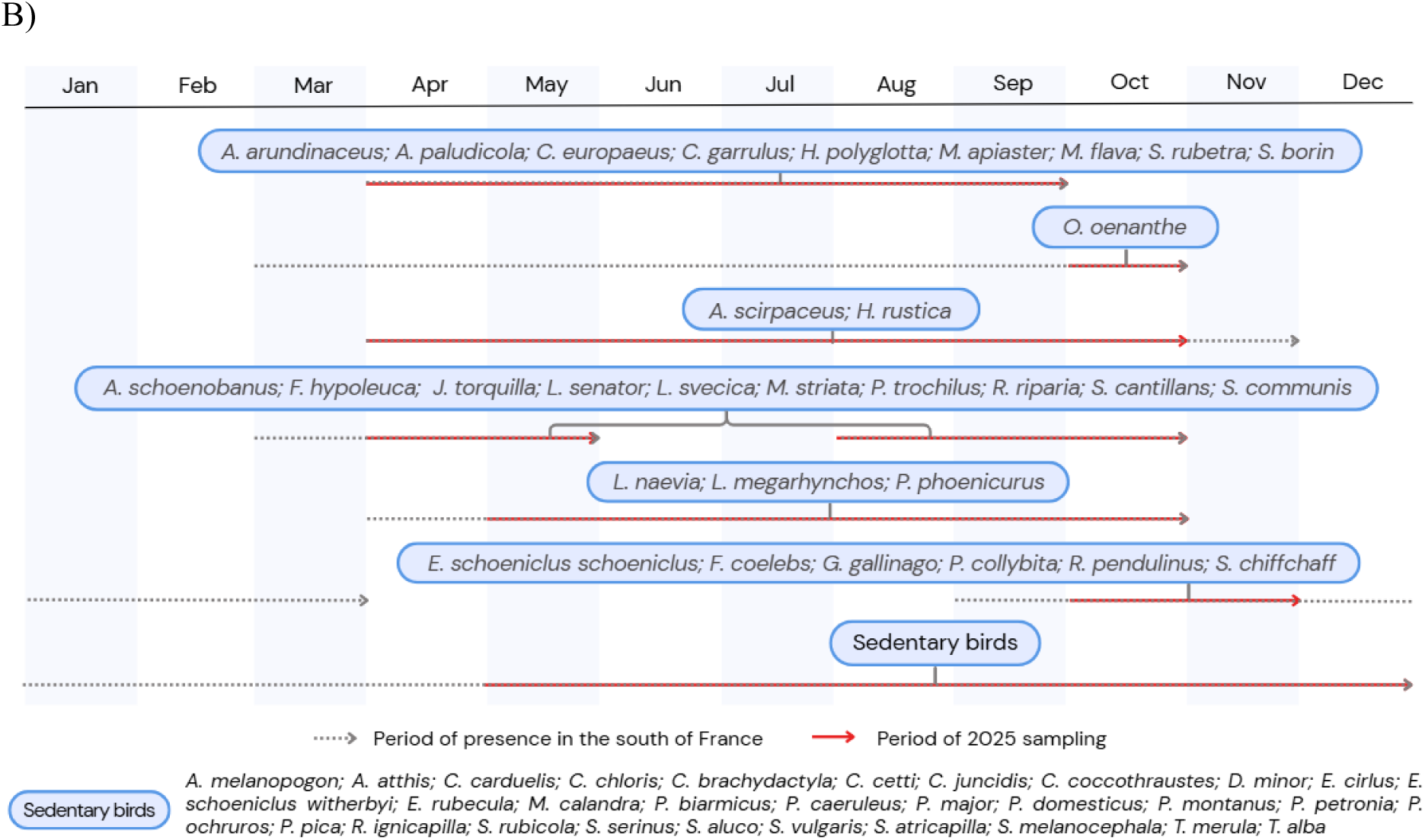
Annual chronological timelines of the sampling periods for bird species collected in 2024 (A) and 2025 (B), based on their periods of presence in the Camargue region.

### Circulation of USUV/WNV in avifauna

The distribution of USUV and WNV detections across all sampled avifauna, considering 2024 and 2025 together, showed a significant difference in detection frequencies between the two viruses (McNemar’s test, p < 0.001). The overall prevalence was higher for USUV, at 11.9% (303/2552; 95% CI: 10.7–13.2), than for WNV, at 4.3% (110/2552; 95% CI: 3.6–5.2), indicating a clear dominance of USUV detections. Differences in virus circulation between migratory (Figure 4A) and sedentary (Figure 4B) bird species were observed for WNV (χ² test, p = 0.006), but not for USUV (χ² test, p = 0.08). Overall, USUV prevalence was 13.2% (140/1058; 95% CI: 11.3–15.4) in sedentary birds and 10.9% (163/1494; 95% CI: 9.4–12.6) in migratory birds, whereas WNV prevalence was 5.7% (60/1058; 95% CI: 4.4–7.2) and 3.3% (50/1494; 95% CI: 2.5–4.3), respectively. Across both ecological groups, detection frequencies differed significantly between USUV and WNV (McNemar’s test, p < 0.001). Given the higher prevalence estimates for USUV compared with WNV, these results indicate a predominance of USUV detections in both sedentary and migratory birds. At the species level, relatively high prevalence values were observed in several sedentary species such as great tit (*Parus major*) (USUV: 37.3%, 25/67; WNV: 13.6%, 8/67) and house sparrow (*Passer domesticus*) (USUV: 27.7%, 28/101; WNV: 16.8%, 17/101). Among migratory species, detections were largely driven by highly sampled species such as barn swallow (*Hirundo rustica*) (USUV: 12.2%, 25/205), while evidence of exposure to both viruses was observed in species such as European bee-eater (*Merops apiaster*) (USUV: 19.6%, 9/46; WNV: 26.1%, 12/46). At the species level, prevalence estimates varied widely within both groups, reflecting heterogeneous exposure patterns as well as the influence of sample size, with higher values often observed in species with limited numbers of individuals. Overall, these results indicate that virus circulation occurred in both ecological groups but tended to be higher in sedentary species, whereas migratory birds appeared to contribute less to the overall viral burden.

**Figure 4:**
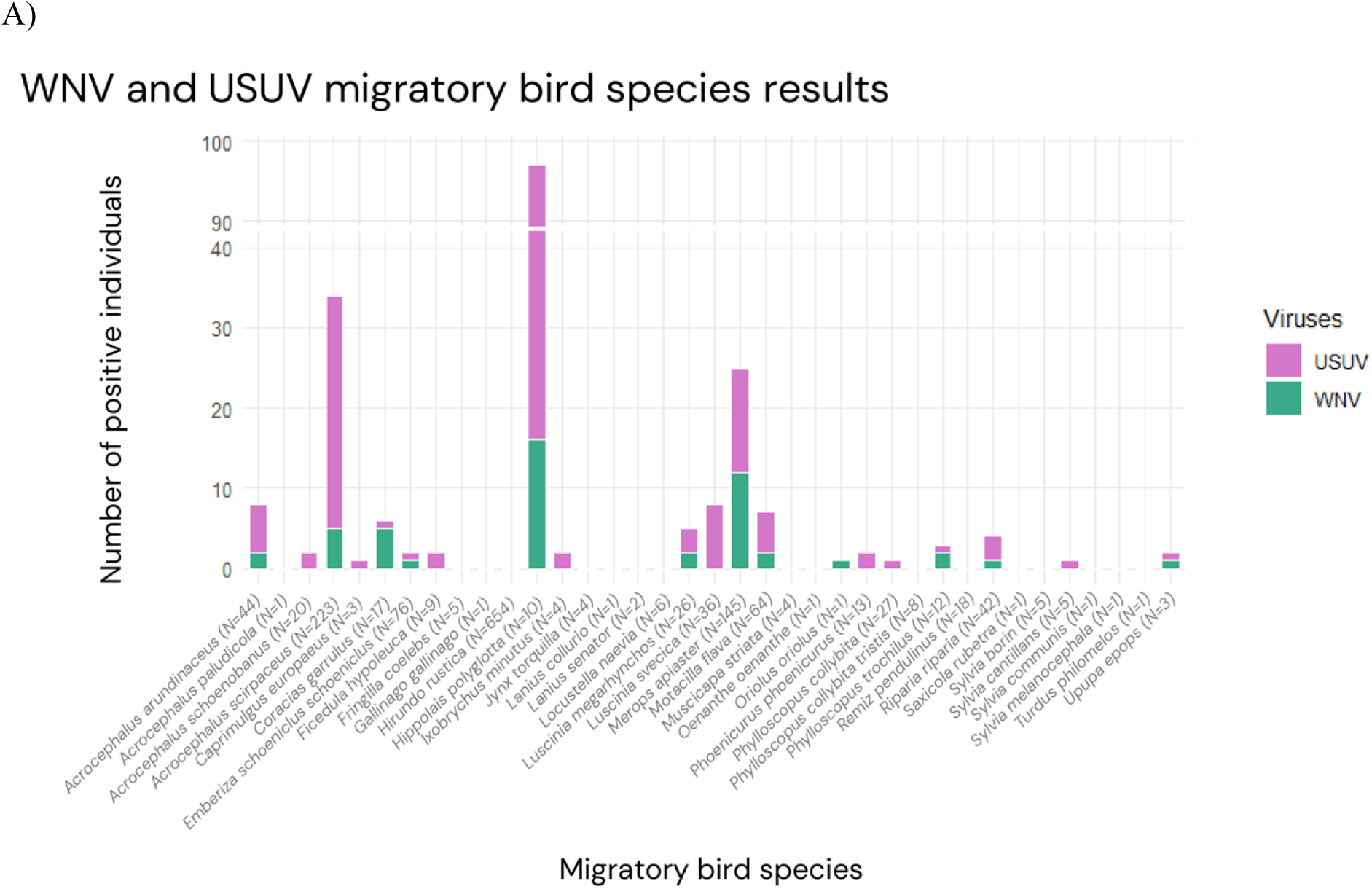
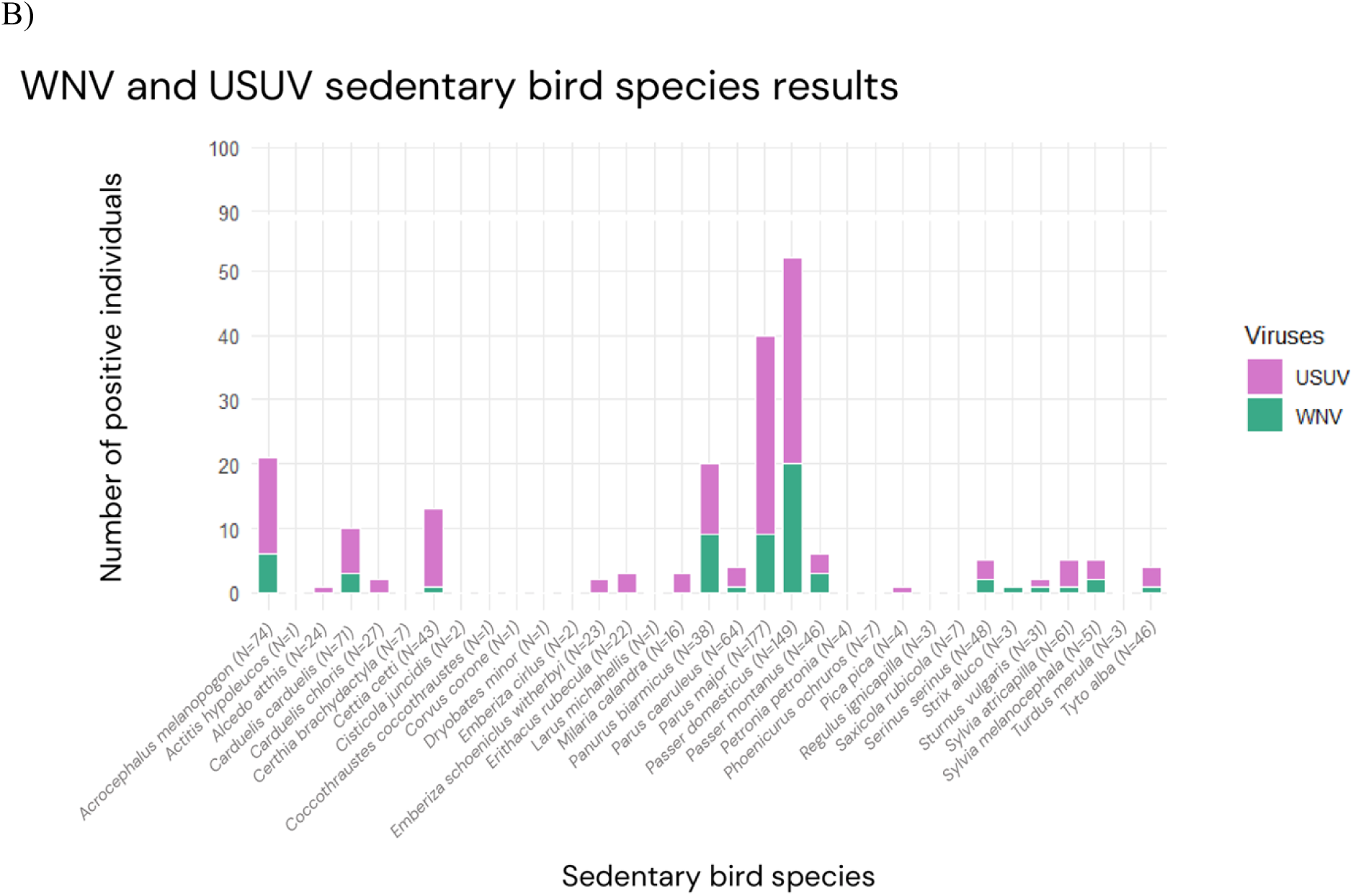
WNV and USUV positive detections by bird species in 2024 and 2025, stratified by migratory behavior. Number of individuals testing positive for West Nile virus (green) and Usutu virus (purple) among migratory (A) and sedentary (B) bird species on both years. Sample sizes for each species are indicated in parentheses along the x-axis.

At the order level, most sampled individuals belonged to Passeriformes (2299/2552), compared to 253 individuals from other avian orders (Supplementary Table 1). Consequently, this order accounted for a large proportion of virus detections, representing 92.7% of USUV-positive individuals (281/303) and 81.8% of WNV-positive individuals (90/110). However, no significant association between Passeriformes and USUV positivity was observed (χ² test, p = 0.16). In contrast, WNV positivity differed significantly between Passeriformes and other avian orders (χ² test, p = 0.003), with proportionally more positives detected among non-passeriform individuals in our dataset. These results should nevertheless be interpreted with caution due to the strong imbalance in sampling effort between taxonomic groups and the ecological heterogeneity across species.

Across both years, co-infections with USUV and WNV remained uncommon, accounting for 2.0% of all sampled birds (51/2552; 95% CI: 1.5–2.6), and were detected in a limited set of host species, with marked heterogeneity in prevalence across species. Co-infections were markedly more frequent in 2024, with 42 double-positive individuals (4.1%; 42/1032), than in 2025, when only 9 cases were detected (0.6%; 9/1520) (Fisher’s exact test, p < 0.001). When considering the cumulative dataset, co-infection prevalence ranged from less than 1% in highly sampled species, such as the barn swallow (0.8%, 5/654), to values exceeding 10% in species with smaller sample sizes, including the bearded reedling (10.5%, 4/38) and the Eurasian tree sparrow (6.5%, 3/46). Among common sedentary species, co-infections were observed in the house sparrow (8.1%, 12/149) and the great tit (4.0%, 7/177), whereas migratory species generally exhibited lower co-infection prevalence, with the exception of localized signals in the European bee-eater (2.8%, 4/145) and the European roller (5.9%, 1/17). Several other species showed sporadic co-infections at low prevalence levels (≤4%), reflecting limited but recurrent overlap of USUV and WNV circulation. Taken together, these results suggest that co-infections represent a rare epidemiological outcome, unevenly distributed among host species, and not restricted to a specific ecological guild or taxonomic group.

Host sex was not significantly associated with WNV or USUV positivity. Individuals were classified as male or female, excluding those with undetermined sex (n = 1488). For WNV, positivity did not differ significantly between males (6.1%) and females (4.0%) (Fisher’s exact test, p = 0.17). Similarly, USUV positivity was comparable between males (10.3%) and females (10.4%) (χ² test, p = 1.0). Overall, no significant association was observed between host sex and virus positivity.

Virus positivity differed significantly among age classes for WNV and USUV (χ² tests, p < 0.001 for both analyses). For WNV, the highest positivity rate was observed in *pullus* (PUL; 17.6%), corresponding to the nesting stage characterized by dependence and absence of flight capability, whereas all other age classes showed lower rates, with a maximum of 6.4%. For USUV, volant individuals (VOL; 22.8%), corresponding to the fledgling stage marked by flight ability and increasing autonomy, and PUL (18.3%) exhibited the highest proportions of positive individuals, while older age classes consistently displayed lower positivity. Age was significantly associated with virus positivity, with juvenile stages, particularly *pullus* and volant individuals, showing higher positivity compared to older age classes.

Most captured birds belonged to insectivorous species, either strictly insectivorous (64%) or insectivorous during the breeding season and granivorous in winter (30%). No significant association was detected between dietary guild and positivity for either WNV or USUV (USUV: χ² test, p = 0.12; WNV: Fisher’s exact test, p = 0.07). Dietary specialization was not associated with virus positivity, suggesting that feeding ecology alone does not explain differences in infection risk within the sampled avian communities.

### Temporal Dynamics and Spatial Distribution of USUV and WNV

The distribution of USUV and WNV detections revealed significant inter-annual differences in virus circulation (USUV: χ² test, p = 0.007; WNV: χ² test, p < 0.001). In 2024, the overall prevalence reached 14.0% for USUV (144/1032; 95% CI: 11.9–16.0) and 7.7% for WNV (79/1032; 95% CI: 6.0–9.3). In 2025, lower prevalence levels were observed, with 10.5% for USUV (159/1520; 95% CI: 8.9–12.0) and 2.0% for WNV (31/1520; 95% CI: 1.3–2.8). This shift was characterized by a marked reduction in WNV detections and a more moderate decrease in USUV prevalence, while USUV remained the most frequently detected virus in both years.

The monthly distribution of USUV and WNV detections revealed distinct patterns (Fisher’s exact test, p < 0.001). In 2024 (Figure 5A; Table 1A), viral activity was concentrated between May and October, with a pronounced synchronous peak in June (WNV: 64 positives, 54.2%; USUV: 54 positives, 45.8%), representing the highest level of detection for both viruses. Initial detections occurred in May, corresponding to the first month of sampling in 2024 (WNV: 5, 16.7%; USUV: 6, 20.0%). In July, WNV detections declined markedly (6 positives, 5.5%), while USUV remained relatively elevated (31 positives, 28.4%). From August to October, USUV persisted at moderate levels (18 positives per month; 5.8–9.8%), whereas WNV circulation remained sporadic (≤3 positives per month, ≤1.0%). No viral detections were recorded in November–December for WNV, and USUV dropped to minimal levels (1 case in November). In 2025 (Figure 5B; Table 1B), viral activity followed a different temporal pattern characterized by an extended transmission window. USUV detections began as early as March (1 positive, 25.0%, based on small sample size) and increased progressively through summer, peaking in September (68 positives, 31.6%), before declining in October (11 positives, 5.0%). In contrast, WNV circulation remained limited throughout the year, with low-level detections from April to October and a modest peak in July (9 positives, 3.5%). Overall, 2024 was characterized by an early-summer synchronous amplification of both viruses, whereas 2025 displayed a prolonged USUV transmission season with a delayed late-summer/early-autumn peak and persistently low WNV activity. Sampling intensity varied across months, with lower numbers of samples tested in early and late periods, resulting in wider confidence intervals for some estimates.

**Figure 5:**
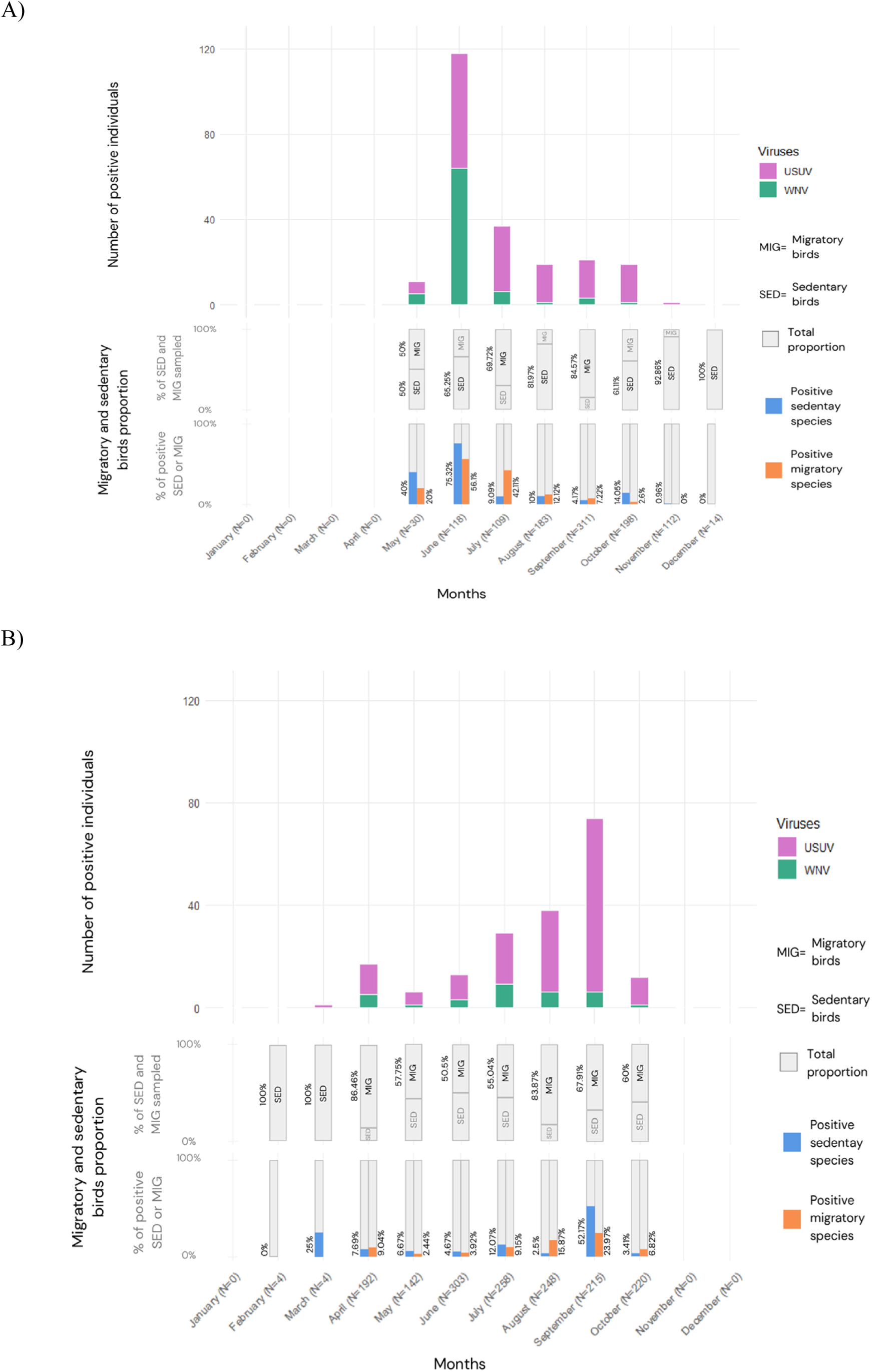
Monthly distribution of WNV and USUV positive birds in 2024 and 2025, proportion of sampled species and proportion of positive species by migratory status. Number of individuals testing positive for WNV (green) and USUV (purple) by month for 2024 (A) and 2025 (B). Sample sizes per month are indicated in parentheses on the x-axis. Proportion of migratory status sampled per month, and proportion of positive by migratory status among the sampled proportion.

**Table 1:** Monthly and locality-based prevalence of West Nile virus (WNV) and Usutu virus (USUV) in 2024 and 2025. Panel A shows the monthly and locality-based prevalence of WNV and USUV in 2024, while Panel B presents the same data for 2025. For each month and locality, the number of positive samples (n), prevalence (%), and 95% confidence interval (95% CI) are reported for both viruses, along with the total number of samples tested.

|  | WNV positive<br>(n) | Prevalence<br>% | 95% CI | USUV positive<br>(n) | Prevalence<br>% | 95% CI | N Total |
| --- | --- | --- | --- | --- | --- | --- | --- |
| **Month of sampling** |  |  |  |  |  |  |  |
| Mai | 5 | 16.7% | 7.3–33.6 | 6 | 20.0% | 9.5–37.3 | 30 |
| Juin | 64 | 54.2% | 45.3–63.0 | 54 | 45.8% | 37.1–54.7 | 118 |
| Juillet | 6 | 5.5% | 2.6–11.5 | 31 | 28.4% | 20.8–37.5 | 109 |
| Aout | 1 | 0.6% | 0.1–3.0 | 18 | 9.8% | 6.3–15.0 | 183 |
| Septembre | 3 | 1.0% | 0.3–2.8 | 18 | 5.8% | 3.7–9.0 | 311 |
| Octobre | 1 | 0.5% | 0.1–2.8 | 18 | 9.1% | 5.8–13.9 | 198 |
| Novembre | 0 | 0.0% | 0.0–3.3 | 1 | 0.9% | 0.2–4.9 | 112 |
| Décembre | 0 | 0.0% | 0.0–21.5 | 0 | 0.0% | 0.0–21.5 | 14 |
| **Locality** |  |  |  |  |  |  |  |
| Aigues-Mortes | 1 | 10.0% | 1.8–40.4 | 0 | 0.0% | 0.0–27.8 | 10 |
| Aigues-Vives | 34 | 8.0% | 5.8–11.0 | 66 | 15.5% | 12.4–19.3 | 425 |
| Montarnaud | 0 | 0.0% | 0.0–22.8 | 1 | 7.7% | 1.4–33.3 | 13 |
| Rouet | 0 | 0.0% | 0.0–65.8 | 0 | 0.0% | 0.0–65.8 | 2 |
| Saint-Gilles | 4 | 80.0% | 37.6–96.4 | 1 | 20.0% | 3.6–62.5 | 5 |
| Saint-Laurent-D’Aigouze | 22 | 4.2% | 2.8–6.2 | 58 | 11.0% | 8.6–14.0 | 527 |
| Vauvert | 18 | 36.0% | 24.1–49.9 | 18 | 36.0% | 24.1–49.9 | 50 |

**Table 1:** Monthly and locality-based prevalence of West Nile virus (WNV) and Usutu virus (USUV) in 2024 and 2025.
|  | WNV positive<br>(n) | Prevalence% | 95% CI | USUV positive<br>(n) | Prevalence% | 95% CI | N Total |
| --- | --- | --- | --- | --- | --- | --- | --- |
| Month of sampling |  |  |  |  |  |  |  |
| Février | 0 | 0.0% | 0.0–49.0 | 0 | 0.0% | 0.0–49.0 | 4 |
| Mars | 0 | 0.0% | 0.0–49.0 | 1 | 25.0% | 4.6–70.0 | 4 |
| Avril | 5 | 2.6% | 1.1–6.0 | 12 | 6.3% | 3.1–10.6 | 192 |
| Mai | 1 | 0.7% | 0.1–3.9 | 5 | 3.5% | 1.5–8.0 | 142 |
| Juin | 3 | 1.0% | 0.3–2.9 | 10 | 3.3% | 1.8–6.0 | 303 |
| Juillet | 9 | 3.5% | 1.9–6.5 | 20 | 7.8% | 5.1–11.7 | 258 |
| Aout | 6 | 2.4% | 1.1–5.2 | 32 | 12.9% | 9.3–17.7 | 248 |
| Septembre | 6 | 2.8% | 1.3–6.0 | 68 | 31.6% | 25.8–38.1 | 215 |
| Octobre | 1 | 0.5% | 0.1–2.5 | 11 | 5.0% | 2.8–8.7 | 220 |
| Locality |  |  |  |  |  |  |  |
| Agde | 0 | 0.0% | 0.0–3.2 | 3 | 2.6% | 0.9–7.3 | 116 |
| Aigues-Mortes | 1 | 6.3% | 1.1–28.3 | 2 | 12.5% | 3.5–36.0 | 16 |
| Aigues-Vives | 3 | 1.1% | 0.4–3.2 | 9 | 3.3% | 1.8–6.2 | 272 |
| Arles | 0 | 0.0% | 0.0–56.2 | 0 | 0.0% | 0.0–56.2 | 3 |
| Codognan | 0 | 0.0% | 0.0–79.4 | 0 | 0.0% | 0.0–79.4 | 1 |
| Le Cailar | 0 | 0.0% | 0.0–27.8 | 1 | 10.0% | 1.8–40.4 | 10 |
| Le Grau-du-Roi | 0 | 0.0% | 0.0–35.4 | 0 | 0.0% | 0.0–35.4 | 7 |
| Montarnaud | 0 | 0.0% | 0.0–49.0 | 1 | 25.0% | 4.6–69.9 | 4 |
| Montpellier | 1 | 1.2% | 0.2–6.5 | 6 | 7.2% | 3.4–14.9 | 83 |
| Murviel-lès-Montpellier | 0 | 0.0% | 0.0–65.8 | 0 | 0.0% | 0.0–65.8 | 2 |
| Rouet | 1 | 1.1% | 0.2–5.8 | 11 | 11.8% | 6.7–20.0 | 93 |
| Saint-Gilles | 7 | 6.5% | 3.2–12.8 | 5 | 4.6% | 2.0–10.4 | 108 |
| Saint-Laurent-d’Aigouze | 7 | 1.9% | 0.9–3.9 | 74 | 20.1% | 16.3–24.5 | 368 |
| Saintes-Maries-de-la-Mer | 0 | 0.0% | 0.0–39.0 | 0 | 0.0% | 0.0–39.0 | 6 |
| Vauvert | 4 | 2.3% | 0.9–5.7 | 28 | 15.9% | 11.2–22.0 | 176 |
| Villeneuve-lès-Maguelone | 7 | 2.8% | 1.3–5.6 | 19 | 7.5% | 4.8–11.3 | 255 |

Spatially, viral circulation displayed a heterogeneous distribution across sampling sites, with differences observed between 2024 and 2025. In 2024 (Figure 6A; Table 1A), WNV detections were recorded across several localities, including rural areas (Saint-Gilles, Vauvert, Saint-Laurent-d’Aigouze) and a suburban area (Aigues-Vives); however, site-specific proportions should be interpreted with caution due to limited sample sizes in some locations. In contrast, USUV detections were more widely distributed, with higher proportions observed in rural areas such as Vauvert (36.0%) and Saint-Laurent-d’Aigouze (11.0%), and in the suburban area of Aigues-Vives (15.5%), providing more robust site-level estimates. In 2025 (Figure 6B; Table 1B), USUV activity remained substantial across several rural sites, including Saint-Laurent-d’Aigouze (20.1%), Vauvert (15.9%), and Rouet (11.8%). In contrast, WNV prevalence remained low across most sites, with slightly higher values observed in Saint-Gilles (6.5%) and Villeneuve-lès-Maguelone (2.8%). Notably, a small number of USUV-positive birds (n = 6) and a single WNV-positive bird were also detected in 2025 within the city of Montpellier, the only urban site, indicating the presence of viral circulation in urban environments. Across both years combined, the highest proportion of positive cases was observed in rural areas (17.5%; 396/2268), followed by urban (8.4%; 7/83) and peri-urban areas (5.5%; 10/182), indicating a significant association between locality type and virus detection (χ² test, p = 0.002 for USUV; Fisher’s exact test, p = 0.02 for WNV).

**Figure 6:**
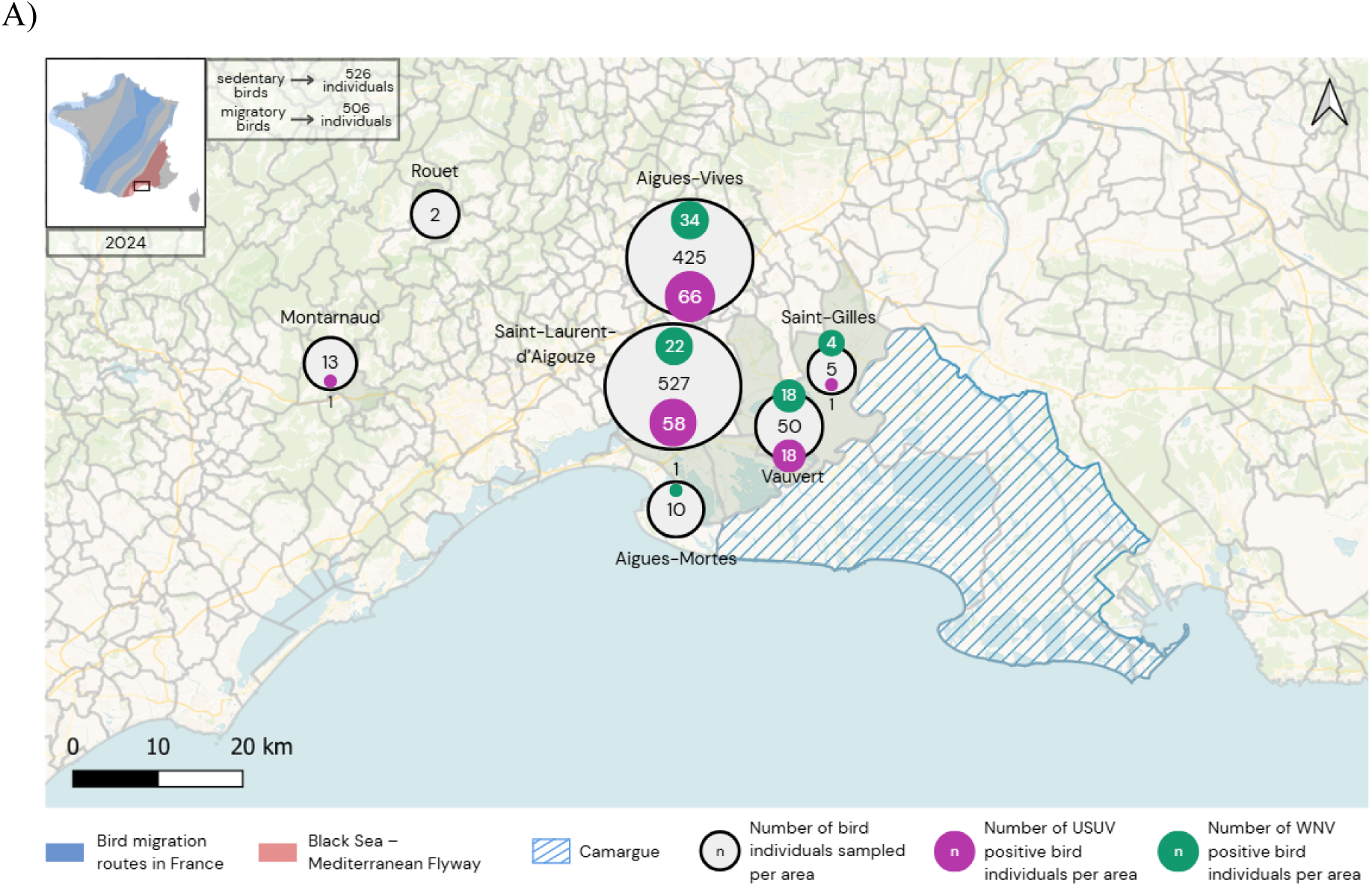
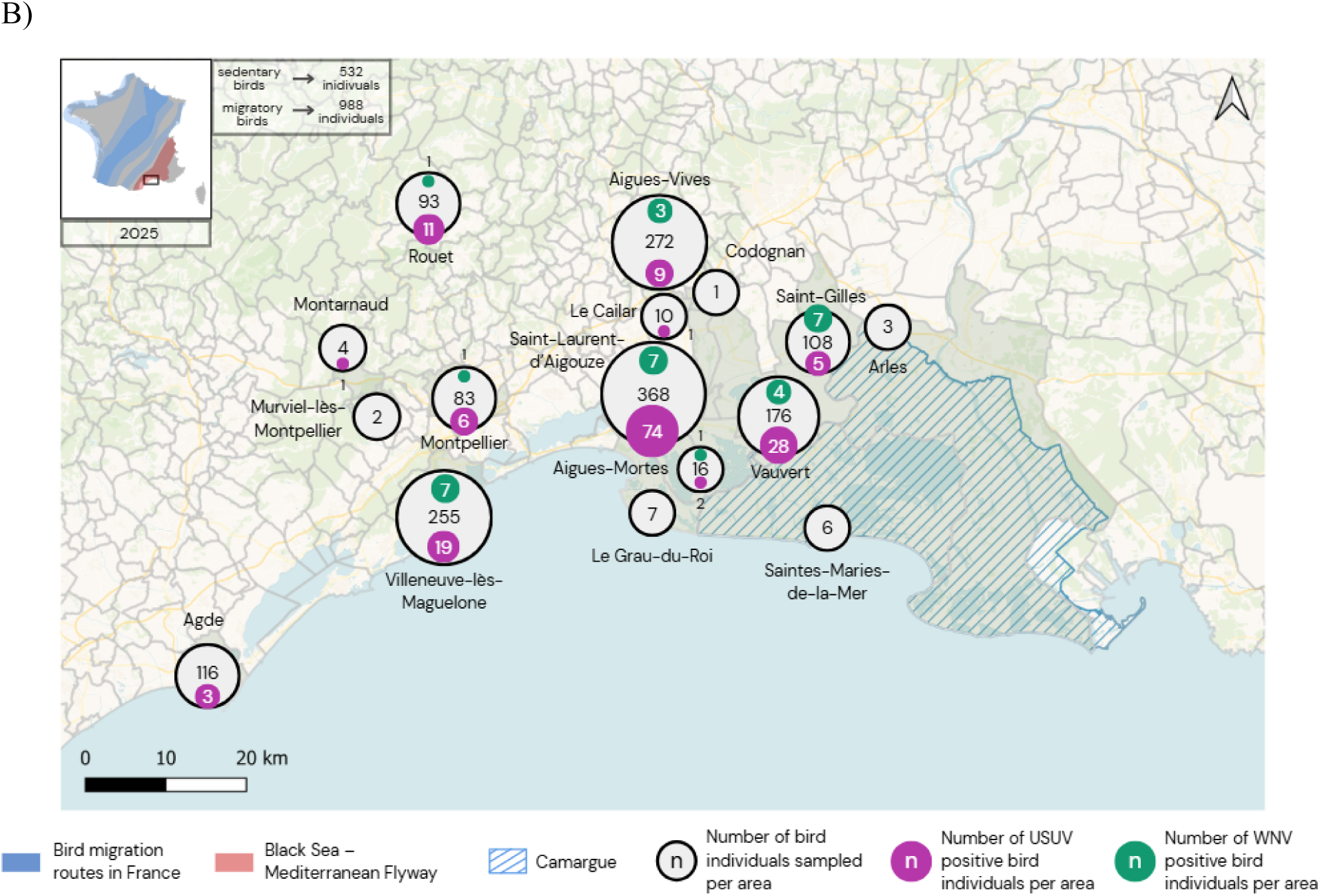
Spatial distribution of USUV and WNV infections in birds captured in 2024 (A) and 2025 (B). Sampling sites are shown with the number of birds testing positive for USUV (purple circles) and WNV (green circles). The total number of captures at each site is indicated within the corresponding grey circle. Hatched areas represent wetlands of the Camargue Park. Insets show the total numbers of sedentary and migratory birds captured across all sites each year.

Separate mixed-effects logistic regression models were fitted for USUV and WNV detection. After model selection, both final multivariable models retained the same explanatory variables: year, season, locality type and age (Table 2), with bird species included as a random effect. Sex, migratory status, Passeriformes status and feeding guild were not retained in either final model. Overall, the multivariable models supported the main patterns observed in the univariable analyses. Detection odds were lower in 2025 than in 2024 for both viruses and higher in rural localities, while age effects were mainly driven by PUL and/or VOL juvenile stages. Seasonal effects were also retained, although the winter effect for USUV should be interpreted cautiously because of the wide confidence interval. In addition, the species random effect accounted for 7.1% of the variance for USUV and 9.9% for WNV, indicating moderate but non-negligible species-level clustering.

**Table 2:**
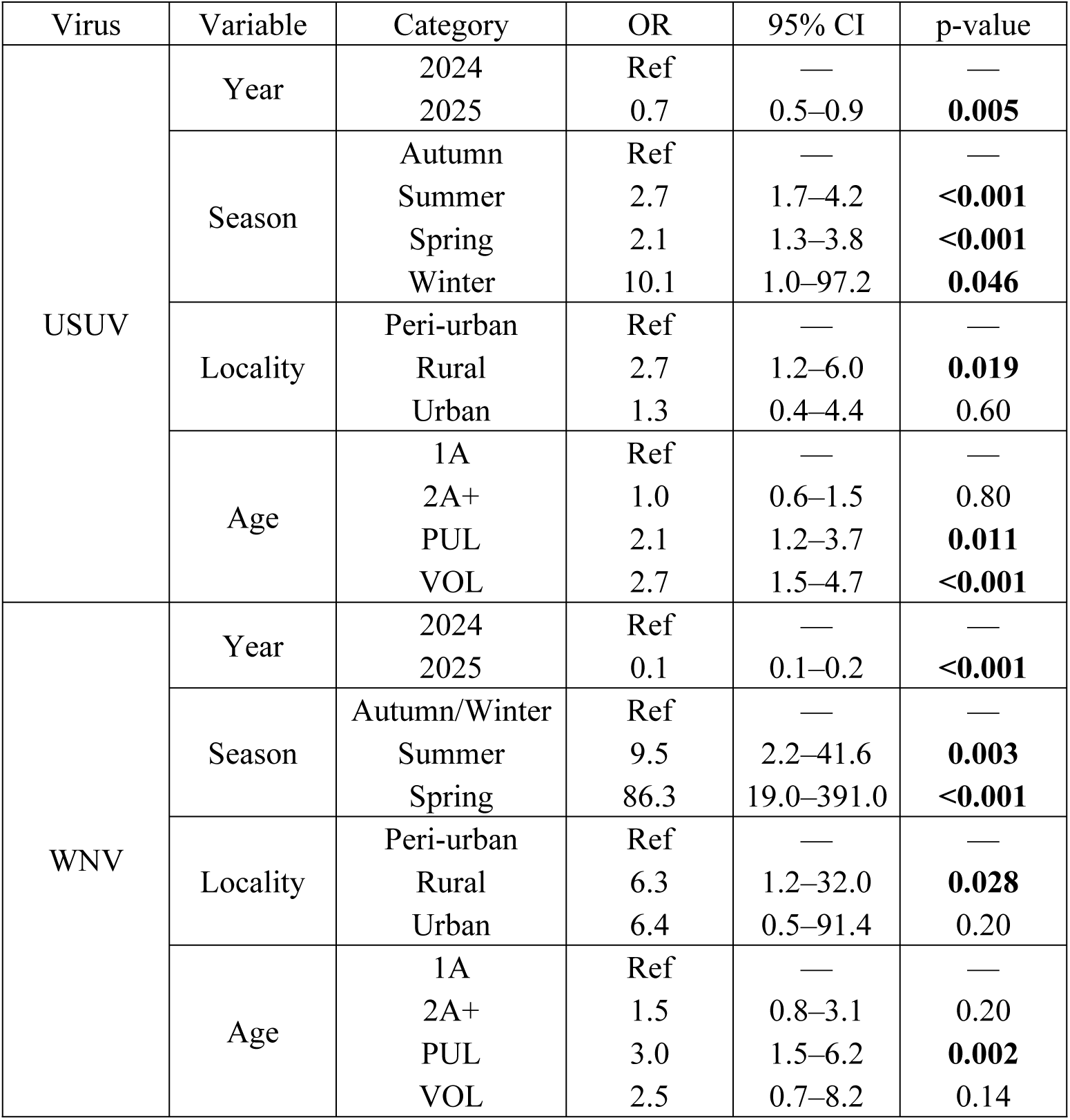
Multivariable mixed-effects logistic regression models for USUV and WNV detection in wild birds. Results are presented as odds ratios (OR), 95% confidence intervals (95% CI) and p-values. Values in bold indicate statistically significant associations at p < 0.05. Reference categories are indicated as “Ref.”. For the WNV model, autumn and winter were combined as the reference category because of the low number of winter observations.

A total of 93 birds were captured on at least two occasions during the study period, allowing a longitudinal assessment of virological status at discrete capture dates (Figure 7). Although most individuals were tested negative at all capture dates, individual trajectories revealed temporal heterogeneity. Among birds that tested positive at least once, most corresponded either to individuals that were already positive at their first capture and subsequently became negative, for whom the onset of positivity could not be determined, or to individuals that were initially negative and became positive at a later capture, allowing in this case the identification of a bounded window of infection acquisition but without information on the timing of negativation. Three individuals exhibited a negative–positive–negative qPCR sequence. For these individuals, the maximum duration of infection was conservatively estimated as the interval between the last negative test preceding positivity and the first negative test following positivity. These intervals ranged from 24 to 41 days, providing upper bounds for the duration of infection given the discrete temporal resolution of the sampling. One individual exhibited two consecutive positive qPCR results for USUV across captures separated by four days. Additional sequential sampling would be necessary to refine the duration of detection according to species. No individual showed repeated positive detections over longer time intervals, and no evidence of prolonged or long-term persistent qPCR positivity was observed at the temporal resolution of the sampling.

**Figure 7:**
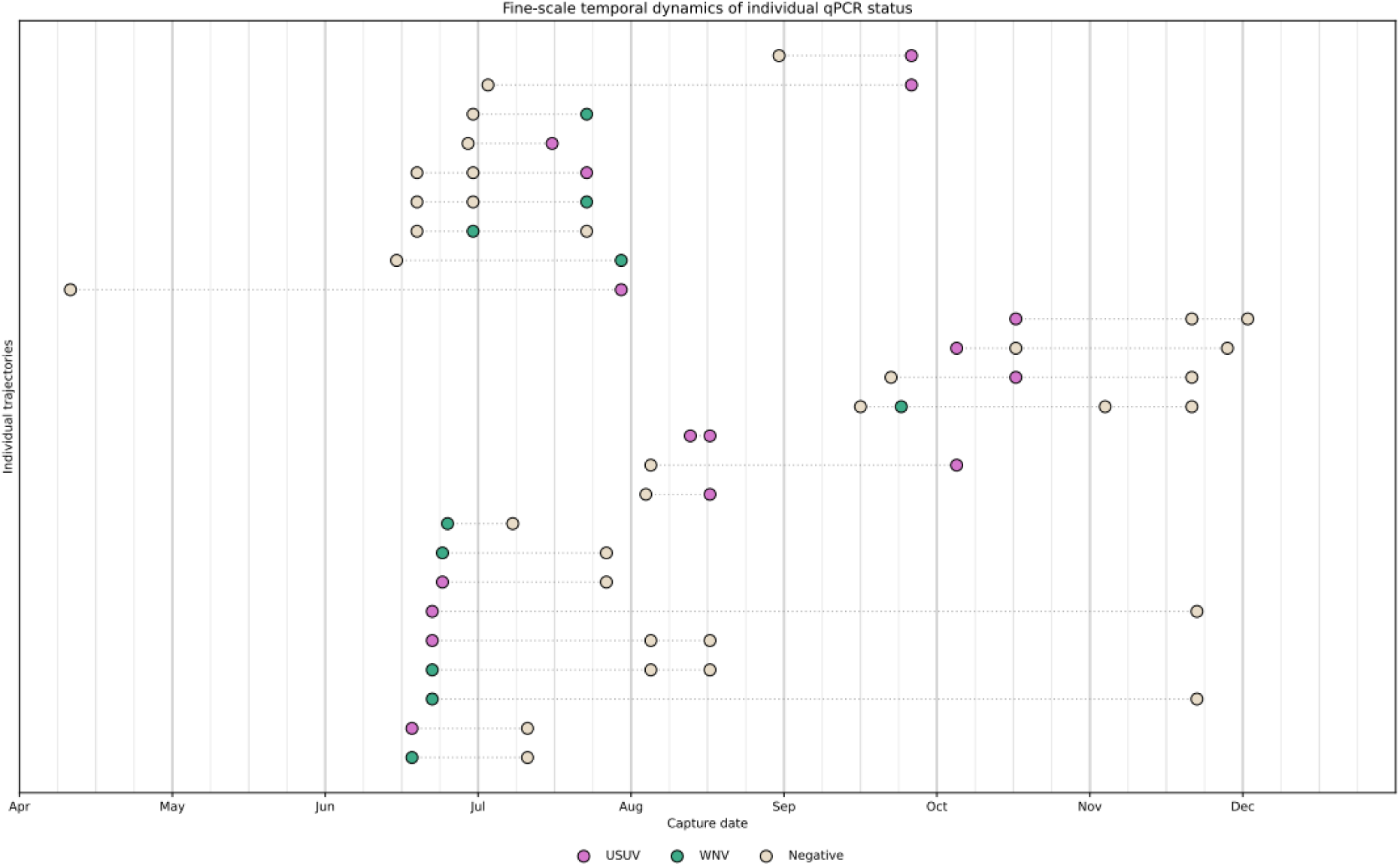
Fine-scale temporal dynamics of individual infection status in 2024 and 2025. Each point represents the positivity status observed at a given capture date for a single individual. Colors indicate viral status (USUV, WNV, or negative). Dashed lines connect successive captures of the same individual and indicate follow-up continuity only, without inferring infection status between capture events. Vertical background lines indicate calendar months (bold), mid-months (medium), and quarter-months (light).

### Partial sequencing and phylogenetic characterization of WNV and USUV

We were unable to obtain complete genome sequences from our samples due to the limited amount of material available from swabs or droppings. Nevertheless, several partial sequences were successfully generated, confirming sample positivity (Figure 8). Sequence analysis of avian samples indicated the circulation of WNV lineage 1 in 2024, from which sequences were successfully obtained. In contrast, mosquito analyses conducted in 2025 revealed the presence of lineage 2 in our study area (43). For USUV, phylogenetic analysis showed that the strains detected in France belonged to lineage Europe 1, a lineage historically circulating mainly in southern and eastern Europe (Italy, Austria, Hungary, Serbia).

**Figure 8:**
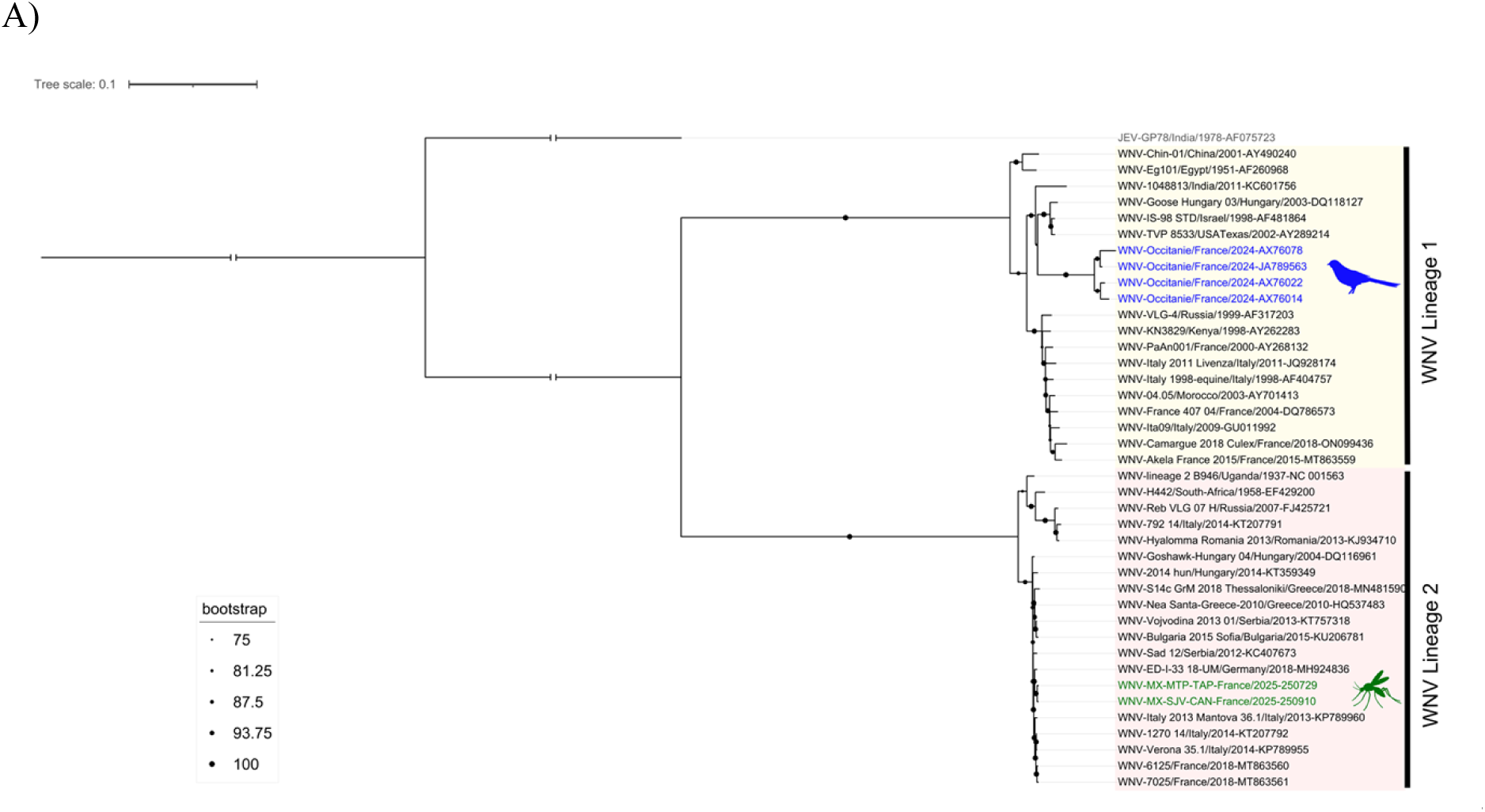
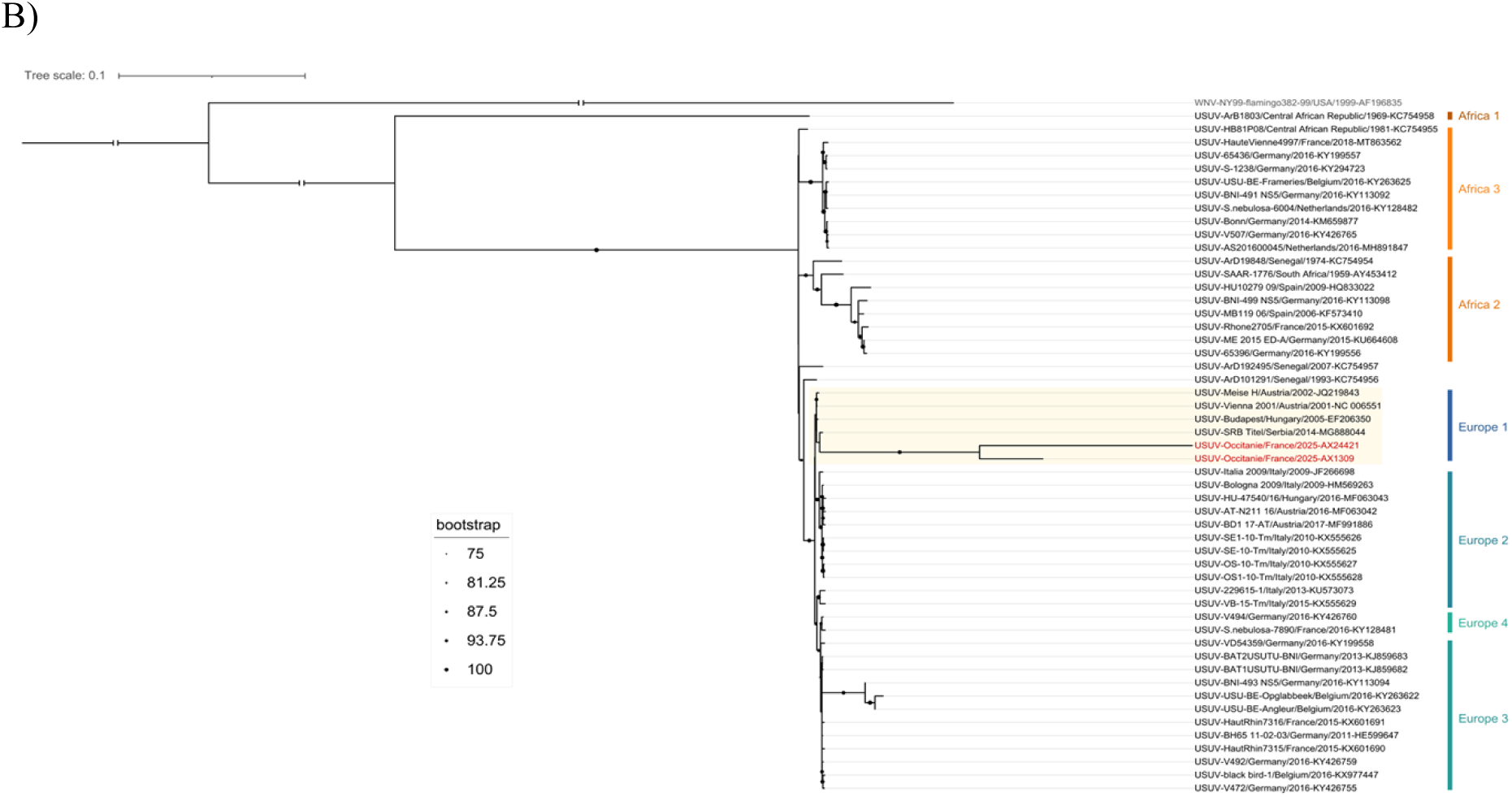
Phylogeny of representative sequences of WNV (A) and USUV (B). (A) Tree was rooted with a sequence of Japanese encephalitis virus (JEV) in grey. Sequences generated for this study are shown in bold green for samples from mosquitoes and in bold blue for avian samples. Grey squares on branch indicate bootstrap score ≥75%. Scale bar represents nucleotide substitutions per site. (B) Phylogeny of representative sequences of Usutu Virus (USUV). Tree was rooted with a sequence of West Nile Virus (WNV) in grey. Sequences generated for this study are shown in bold red. Black points on branch indicate bootstrap score ≥75%. Scale bar represents nucleotide substitutions per site.

### Birds as Early Sentinels of WNV and USUV Circulation: Seasonal Dynamics in 2024–2025

In both years, avian sampling provided the earliest detection of WNV and USUV circulation compared to vectors or dead-end hosts, with first positives occurring several weeks before detections in mosquito pools, equine cases, or human infections (Figure 9). In 2024, initial avian detections occurred in late May, preceding the first human WNV cases (early July), mosquito positives (early August) and equine cases (early September), with viral detections in birds extending into late October–early November. The 2025 season showed earlier onset with USUV-positive birds detected in late March, and WNV in avian samples and mosquito samples in late April, followed by the first WNV human case early July and WNV equine case in early August.

**Figure 9:**
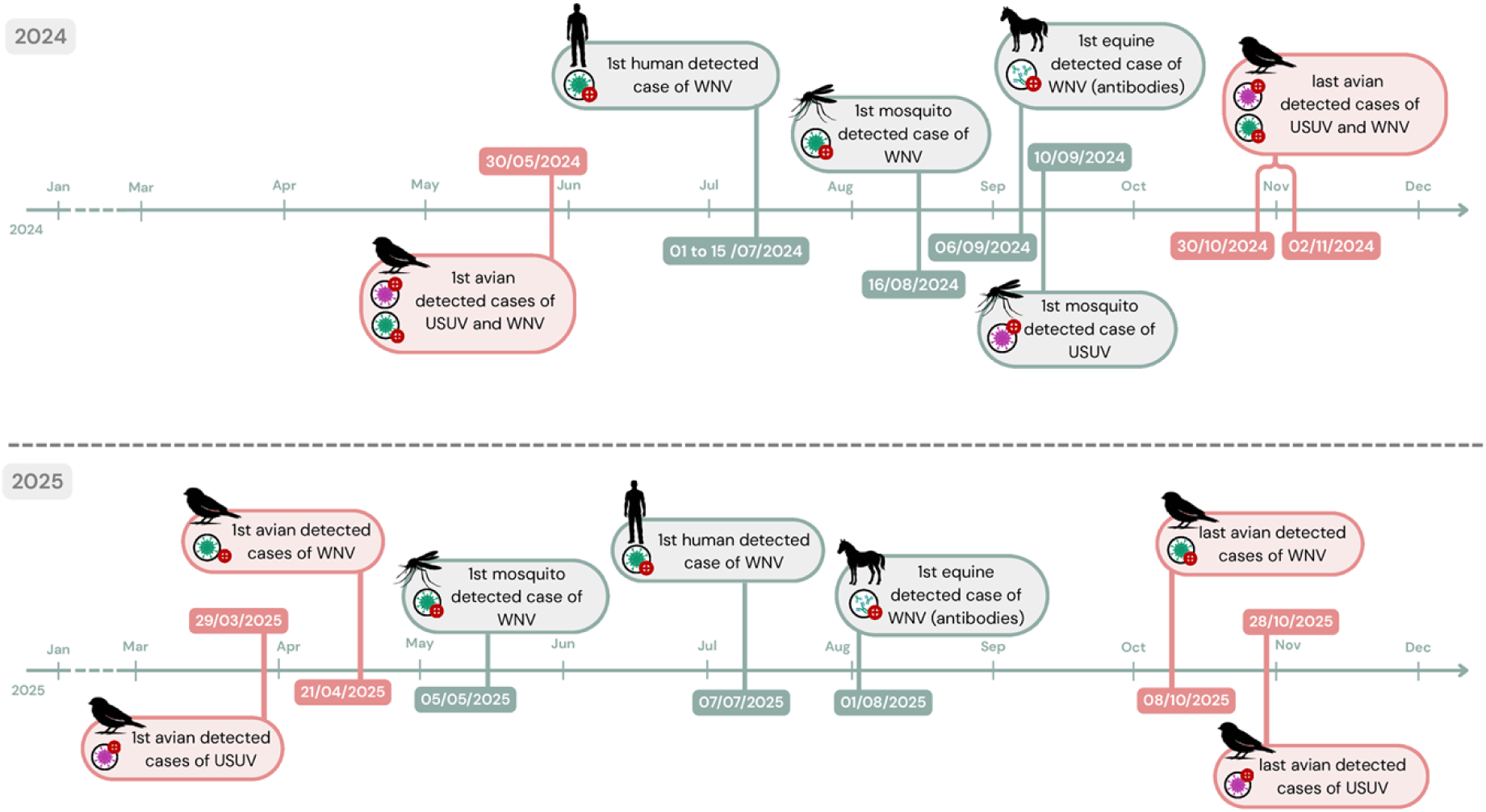
Temporal dynamics of WNV and USUV circulation across multiple surveillance matrices in 2024 and 2025 in Occitanie. Timeline showing first and last detections of WNV (green) and USUV (purple) in birds (orange), and only first detections of WNV and USUV in mosquitoes, equines, and humans (blue). All bird data were generated in this study, whereas data on human cases were obtained from Santé Publique France, equine data from RESPE, and mosquito data from (44).

## Discussion

Our two-year avian molecular surveillance in the Camargue wetlands and urban and peri-urban areas of Montpellier (Occitanie, France) revealed sustained circulation of USUV and WNV. USUV detections were significantly more frequent than WNV detections, with overall prevalences of 11.9% and 4.3%, respectively, and this predominance was observed in both sedentary and migratory bird species. Passeriformes represented the majority of viral detections in absolute numbers, largely because they constituted most of the sampled individuals (90.1%; 2299/2552). The role of Passeriformes should be interpreted at the species level rather than as a uniform effect of the whole order. Within Passeriformes, several species contributed substantially to viral detections, particularly the great tit and house sparrow among sedentary species, and the barn swallow among migratory species. This species-level pattern is consistent with European studies identifying several passerine species as primary reservoirs for WNV and USUV (45–47). Their abundance, synanthropic behavior, and ecological overlap with ornithophilic *Culex pipiens* may favor frequent mosquito–host contact, as further supported by blood-meal analyses in the Camargue showing substantial feeding on passerines (≈42% of meals vs. ≈18% on waterbirds) (48,49). Depending on the species, communal roosting, ground foraging and close association with human-modified environments may further increase exposure to vectors and help explain their frequent involvement in local virus circulation (48,50–52). Experimental data support this interpretation, indicating that some passerines, such as the house sparrow, can develop viremia levels sufficient to sustain onward viral transmission (53). Although detections outside Passeriformes were limited, several non-passeriform species showed noteworthy patterns. European rollers displayed a relatively high WNV detection rate (29.4%), while European bee-eaters exhibited circulation of both USUV and WNV across a relatively large sample size. These findings suggest that viral circulation was not restricted to passerine hosts and broaden the range of avian hosts documented in this local surveillance context, although such signals should be interpreted with caution given the limited sample sizes for several taxa.

The two consecutive years revealed a marked contrast in virus distribution patterns. In 2024, viral detections were broadly distributed across multiple species and sites, with positives identified in 26 of 54 species across 6 of 7 monitored sites. A pronounced peak occurred in June, temporally aligned with 12 autochthonous human WNV cases reported in Occitanie and 89 confirmed equine cases in mainland France during the 2024 transmission season (15,54), recalling the 2015 Camargue epizootic (55). In 2025, this multi-species pattern shifted toward a more concentrated dynamic, largely driven by a single dominant species, the barn swallow, which was the main contributing species to both USUV and WNV detections. In Occitanie in 2025, 8 human cases and nearly 11 equine cases of West Nile virus infection were reported (56,57). In contrast, most other species showed only limited positivity, with no evidence of widespread amplification comparable to that observed in 2024. Temporal patterns were also strongly structured at the seasonal scale, with higher detection probabilities during periods of vector activity. WNV circulation was mainly observed during spring and summer, whereas USUV was also detected during these seasons but extended into winter, indicating a broader seasonal distribution. This pattern is consistent with recent European-wide syntheses, which increasingly report more widespread and frequent USUV activity than WNV, particularly in central and western Europe and in years with mild autumns (28,45,46,58,59). Mechanistically, vector competence data indicate that local *Culex pipiens* populations can be highly susceptible to USUV, sometimes across broader temperature windows, enabling extended transmission into late autumn under temperate conditions (60,61). Early detection of USUV in March 2025 in a resident bird, the western barn owl (*Tyto alba*), suggests potential low-level persistence through multiple mechanisms: (1) infected *Culex* mosquitoes in diapause, (2) chronic viral shedding in birds (3) fecal–oral transmission, as indicated by the high concordance between cloacal swabs and fecal samples (62–64). Vertical transmission appears negligible (65). These non-vector routes may explain sporadic cold-season outbreaks in confined aviary settings, where transmission in the absence of vectors has been documented (63). Spatially, viral circulation was heterogeneous and significantly associated with locality type, with higher detection probabilities observed in rural environments compared to peri-urban areas, while urban sites showed no significant increase in risk despite evidence of low-level circulation. This pattern is consistent with the distribution of suitable mosquito habitats and host communities, with rural and wetland-associated environments providing favorable conditions for sustained transmission. Site-level observations further support this interpretation, with recurrent detections in several rural localities across both years, whereas urban circulation remained limited. Such interannual and spatio-temporal variability is likely influenced by environmental conditions and vector dynamics: warmer and prolonged autumns can extend mosquito activity, allowing delayed amplification in birds and, subsequently, in humans (66–68). Post-epidemic herd immunity may also have contributed, as the intense viral circulation observed in 2024 could have partially reduced susceptibility in 2025, thereby limiting multi-species amplification. Furthermore, the shift in seasonal infection peaks, from early summer (June–July) in 2024 to later in the season (October) in 2025, is consistent with temporal variability previously documented across Europe (68).

Migratory movements likely facilitate the introduction of the virus, whereas resident species appear to sustain local amplification. Migratory status did not explain a significant proportion of the variation in viral detection in the multivariable model. Nevertheless, beyond these statistical patterns, ecological signals suggest more nuanced processes. Early detections in several migratory species, including *Merops apiaster*, *Luscinia megarhynchos*, and *Upupa epops*, as early as May, indicate that viral circulation may begin with the arrival of migratory birds. This chronology is consistent with previous studies showing that infected migratory birds can transport arboviruses over long distances along flyways (45,46,60). However, a key unresolved question is whether migratory birds arrive already infected or acquire infection locally at stopover or breeding sites. Both pathways are plausible, given (i) phylogeographic and epidemiological evidence for long-distance viral transport and (ii) rapid local exposure where ornithophilic *Culex* overlap with dense bird aggregations (46). Notably, in 2024, the earliest positive detections were observed in European bee-eaters (*Merops apiaster*) in May, shortly after their arrival in the study area (late April–early May), suggesting that infected migratory birds may enter the region already viremic and contribute to seeding local transmission. While local acquisition cannot be excluded, this early detection, preceding peak vector abundance, supports introduction along Mediterranean flyways prior to broader amplification. Conversely, repeated detections observed in several resident species across the transmission season suggest a role in local maintenance of viral circulation. Such patterns are consistent with the widely proposed framework of “introduction by migrants and amplification by residents” described in European WNV and USUV eco-epidemiology (35,45). This interpretation is further supported by the strong contribution of individual species, particularly the barn swallow in 2025, whose high number of detections likely reflects species-level characteristics such as gregarious roosting behavior, prolonged seasonal presence, and co-occurrence with ornithophilic *Culex* in agricultural and peri-urban mosaics, which may transiently shift transmission dominance toward gregarious migratory species at stopover and breeding sites (35). Together, these findings emphasize that virus circulation is better understood at the species level, with migratory movements potentially facilitating virus introduction while resident species contribute to local persistence, in a context-dependent and non-exclusive manner. Disentangling these pathways will require effort-adjusted analysis (e.g., occupancy or hierarchical models), longitudinal ringing/recapture, and targeted sequencing along migration routes to assign infection timing and origin (46).

In contrast to studies reporting demographic invariance in orthoflavivirus infection patterns (69), our results indicate that infection risk is structured by host age, as supported by multivariable analyses. For USUV, both juvenile stages (PUL and VOL) showed significantly higher detection probabilities compared to adults, whereas for WNV, this effect was primarily observed in PUL individuals, with no significant increase detected in VOL birds. These findings highlight the importance of individual-level demographic factors, particularly immunologically naïve cohorts, in shaping transmission dynamics within avian communities. Seasonal pulses of susceptible juveniles may therefore contribute disproportionately to viral amplification. In contrast, sex and dietary guild were not retained in the final multivariable model, suggesting that they do not constitute primary drivers of viral detection when considered alongside other factors. Given the limited subset of individuals for which sex was available, only partial patterns could be assessed. Within this subset, WNV detection appeared higher in males, whereas no clear difference was observed for USUV. These observations should be interpreted cautiously and may reflect sampling constraints rather than a consistent biological effect, although they could also be influenced by sex-specific behavioral patterns, habitat use, or physiological mechanisms affecting susceptibility. Similarly, the absence of a detectable effect of dietary guild is consistent with previous studies (70) and supports the idea that feeding ecology alone plays a limited role compared to habitat use and fine-scale overlap with *Culex* breeding and resting sites. Behavioral shifts, such as post-breeding flock formation and roost relocation to urban parks, riparian zones, and wetlands, likely modulate local host density and mosquito contact, amplifying transmission independently of trophic niche (71).

We documented overlapping transmission of WNV and USUV across the season, with 51 co-infections detected over the two years, totaling 2552 captures (2.0%). The relative rarity of co-infections, despite their occurrence across multiple host species, is consistent with ecological niche overlaps without extensive co-transmission. Co-infections were most frequently observed in species contributing the largest number of detections, particularly house sparrows and great tits, but were also identified in several other species, including barn swallows, European bee-eaters, and bearded reedlings. Interestingly, some species exhibited relatively high prevalence for both viruses but no observed co-infections (e.g., *Upupa epops*), suggesting that co-infection is not solely driven by exposure but may also depend on within-host processes, the timing of infection, or limited sample size, potentially reflecting asynchronous exposure, transient infection windows, or biological interactions limiting co-occurrence at the individual level. These observations indicate broader host overlap rather than confinement to a single ecological group and suggest spatial and temporal niche overlap where *Culex* populations are exposed to both viruses (15,46). Although *in vitro* data indicate that USUV may competitively constrain WNV, the population-level implications of co-infections remain uncertain. Co-circulation of WNV and USUV raises questions regarding lineage-level interactions at the vector–host interface, including competition, facilitation, or cross-immunity. Experimental evidence supports possible antagonistic interactions, as USUV has been shown *in vitro* to inhibit WNV replication via interferon-mediated pathways, potentially contributing to reduced WNV amplification in areas of intense USUV circulation (72). However, field evidence remains limited and sometimes contradictory, and these interactions are likely modulated by ecological drivers such as host availability, temperature regimes, and vector phenology (45,46). From an analytical perspective, the explicit characterization of co-infection distribution in our dataset provides initial insight into shared transmission dynamics. However, extending this approach through dedicated modeling of co-infection patterns, in a framework comparable to that used for single-virus infection, could further disentangle shared and virus-specific drivers of transmission and better quantify the ecological and epidemiological conditions under which co-infections occur. Methodologically, these findings also support the use of multiplex PCR and lineage-aware confirmation (sequencing) to avoid misclassification, detect shifts in viral dominance across seasons, and integrate entomological and avian data within a One Health surveillance framework (15,46,58,59).

Phylogeographic analyses indicate that multiple USUV lineages have entered Europe (33,46,62). In parallel, WNV lineage 2 has progressively replaced lineage 1 across much of Europe and underpins most recent large human and equine outbreaks (43,46). 2024–2025 data show exclusive circulation of WNV lineage 1 in 2024 and a shift to lineage 2 in 2025, contrasting with the historical dominance of lineage 1 in this region and reinforcing evidence for the stable establishment of lineage 2 in southern France, consistent with broader European trends observed since 2010 (15,73,74). For USUV, we identified the Europe 1 lineage, which has been historically circulating in southern and eastern Europe since its emergence in Austria (2001) and Italy (1996) (3,75).

Despite high molecular detection across both years, we observed no conspicuous avian die-offs in the study area. This silent circulation is consistent with several, non-exclusive mechanisms: (i) the infected pool was dominated by tolerant hosts, notably passerines, in which infections are frequently subclinical or mildly symptomatic; (ii) lineage–host interactions likely modulate virulence producing context-dependent outcomes; and (iii) under-ascertainment of carcasses in open wetlands and agricultural mosaics, where detection probability is low and carcasses are rapidly removed by scavengers. This pattern contrasts with documented USUV-associated mortality in blackbirds (*Turdus merula*) and corvids during epizootic years, including severe lesions and high case fatality in experimental infections (with AF3/E3 lineages) (76). Field reports of large blackbird die-offs in several outbreaks highlight strong interspecific differences in pathogenicity and suggest that passive surveillance systems (on dead birds) are biased toward susceptible, conspicuous taxa (corvids, raptors), potentially underestimating burden in passerines where disease is rarely lethal. Consequently, absence of mass mortality should not be interpreted as low transmission risk: subclinical infections in abundant passerines can sustain high viral circulation and raise spillover potential under favorable vector conditions (45).

The multi-matrix surveillance approach (birds, mosquitoes, equines, humans) confirms that avian sampling consistently provides the earliest signal of WNV and USUV circulation, preceding detections in mosquitoes, equines, and humans by 4–12 weeks. In 2024, initial avian positives in late May (none samples before) anticipated human WNV cases by early August and equine neuroinvasive cases by September, while in 2025, birds signaled viral activity as early as late March, months before the first human infection in July. These findings align with decades of evidence from Europe and North America showing that bird-based surveillance, whether through dead bird reporting, sentinel corvids, or targeted sampling offers superior timeliness compared to clinical case detection in incidental hosts (46,77,78). Our data suggest that prevalence thresholds in key passerines (e.g., great tits, house sparrows) could serve as actionable triggers for intensified control strategies, yet such thresholds remain insufficiently standardized across Europe. Integrating avian indicators into One Health frameworks, alongside mosquito and climatic data, has already demonstrated strong predictive performance in regional models and should be scaled to support risk-based, spatially targeted interventions.

Several limitations should be acknowledged. First, molecular detection, as opposed to serology, primarily captures active infections and may therefore underestimate past exposure and cumulative infection risk. Second, uneven sampling effort across species, sites, and seasons may influence detection probabilities and contribute to apparent heterogeneity, particularly for species with small sample sizes. This limitation is especially relevant when interpreting the absence of detection or co-infection in certain taxa. Third, spatial aggregations such as communal roosts may locally inflate infection rates and obscure broader-scale transmission patterns. In addition, although species-level effects were partially accounted for, unmeasured ecological variables (e.g., microhabitat use, vector density, or fine-scale host behavior) may also contribute to the observed variability. Finally, while co-infections were explicitly described, their drivers were not formally modeled, limiting inference on the mechanisms underlying their occurrence.

## Conclusion

Our results demonstrate substantial and early avian circulation of WNV and USUV in southern France, with certain species serving as key maintenance hosts. Unlike passive surveillance, which is often biased toward conspicuous mortality events or clinical cases, active sampling captures a broader range of host species, including those experiencing silent or subclinical infections, thereby providing a more complete picture of viral circulation. Strengthening active bird-based surveillance and integrating it into One Health frameworks will be essential for anticipating outbreaks and guiding timely, targeted public health responses.

## Acknowledgements

We would like to thank the Conservatoire du Littoral, CEN Occitanie and the OFB for the Estagnol and Vagaran/Boulas National Nature Reserve (Villeneuve-les-Maguelone/34), the Ministry of Culture/National Archives for the Espeyran estate (Saint-Gilles/30), the Syndicat Mixte de la Camargue Gardoise for the Tour Carbonnière (Aigues-Mortes/30), the Cougourlier (Saint-Gilles/30), and the Scamandre Discovery Centre (Vauvert/30), Nextstone for the Aigues- Vives quarry (30), and ADENA for the Bagnas National Nature Reserve (Agde/34). The CEFE team thanks the Jardin des Plantes and the Mairie of Montpellier, as well as participants to the capture sessions, in particular Vassily Reach and Samuel Moulin.

## Supplementary

**Supplementary Table 1:**
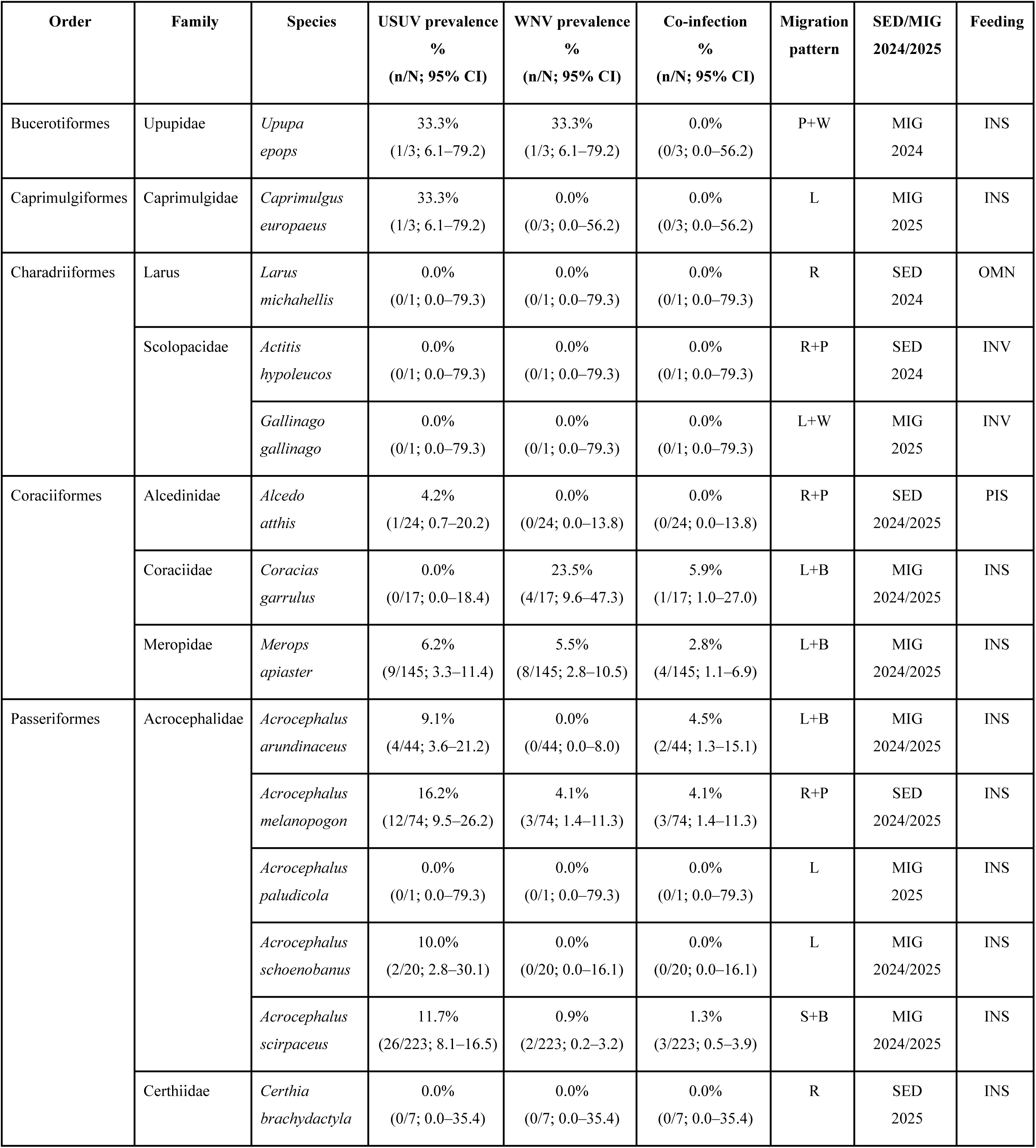
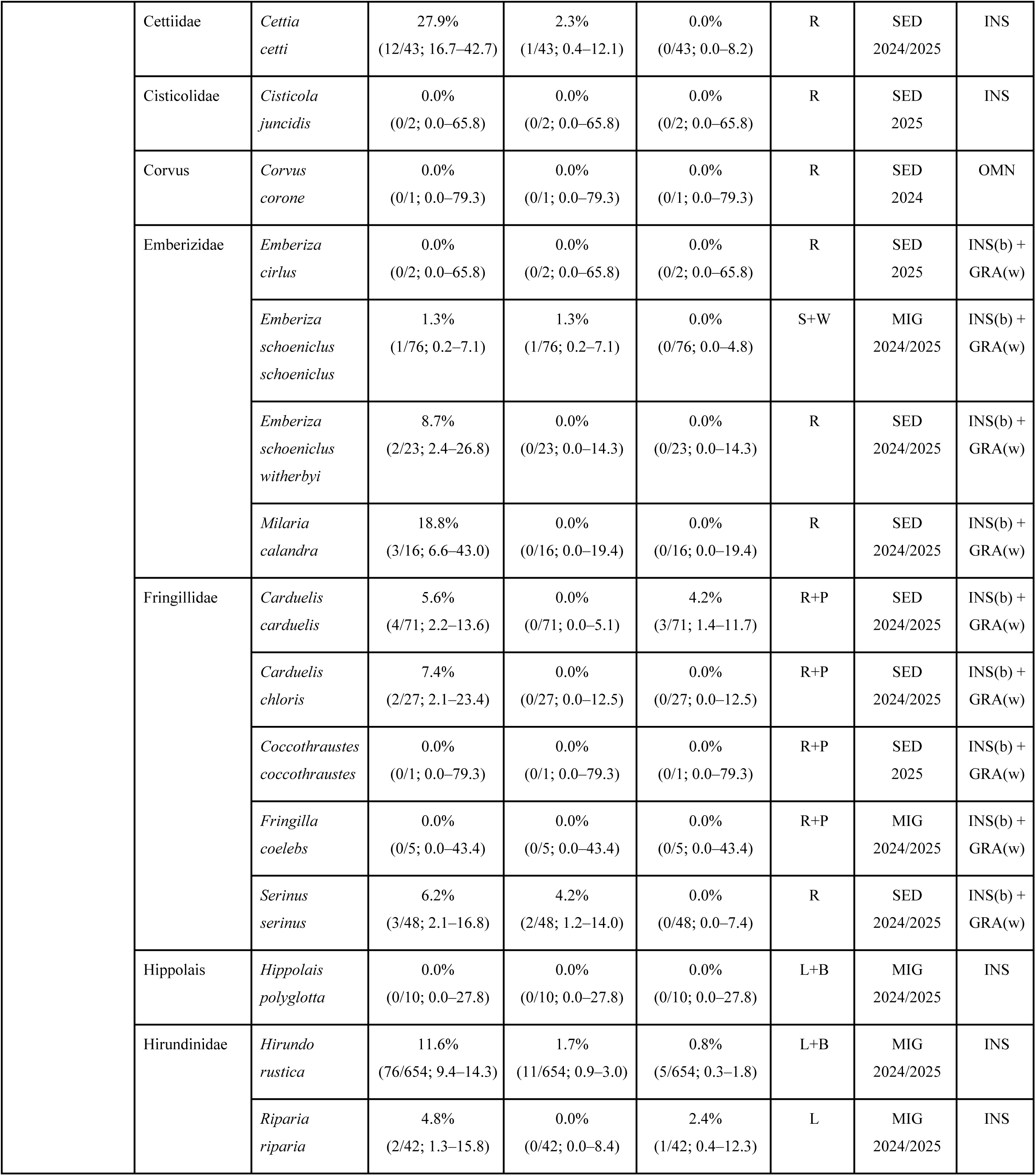
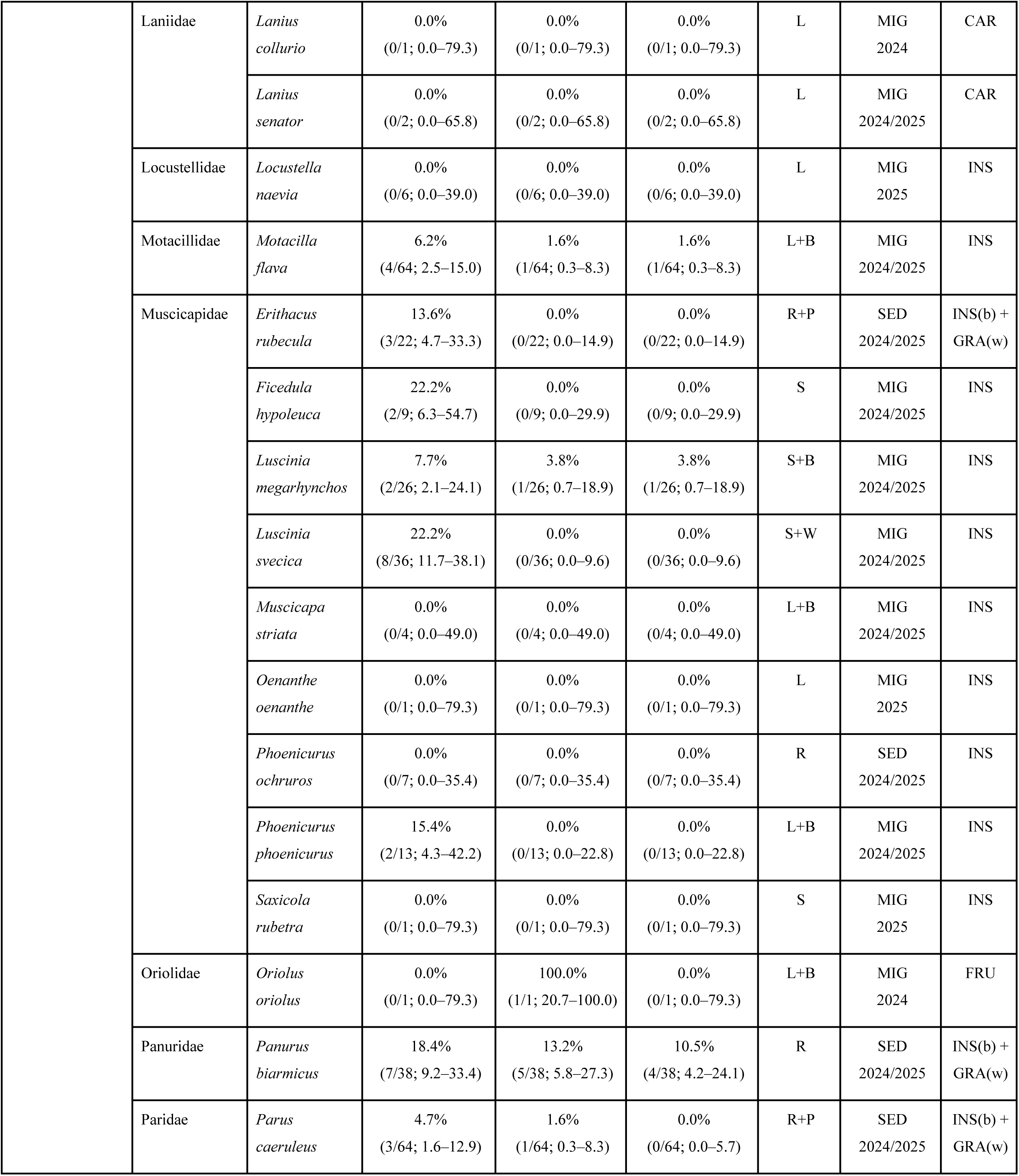
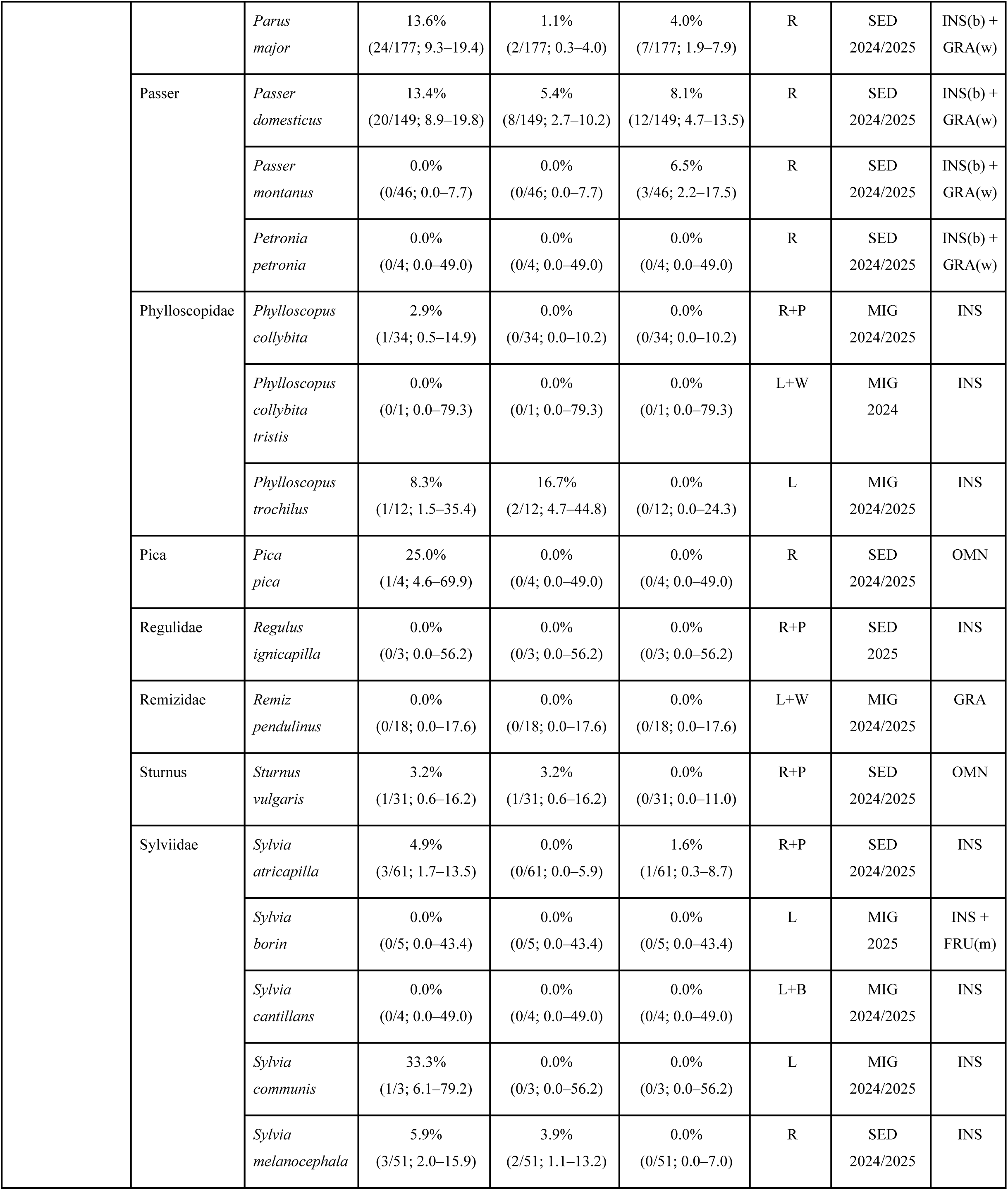
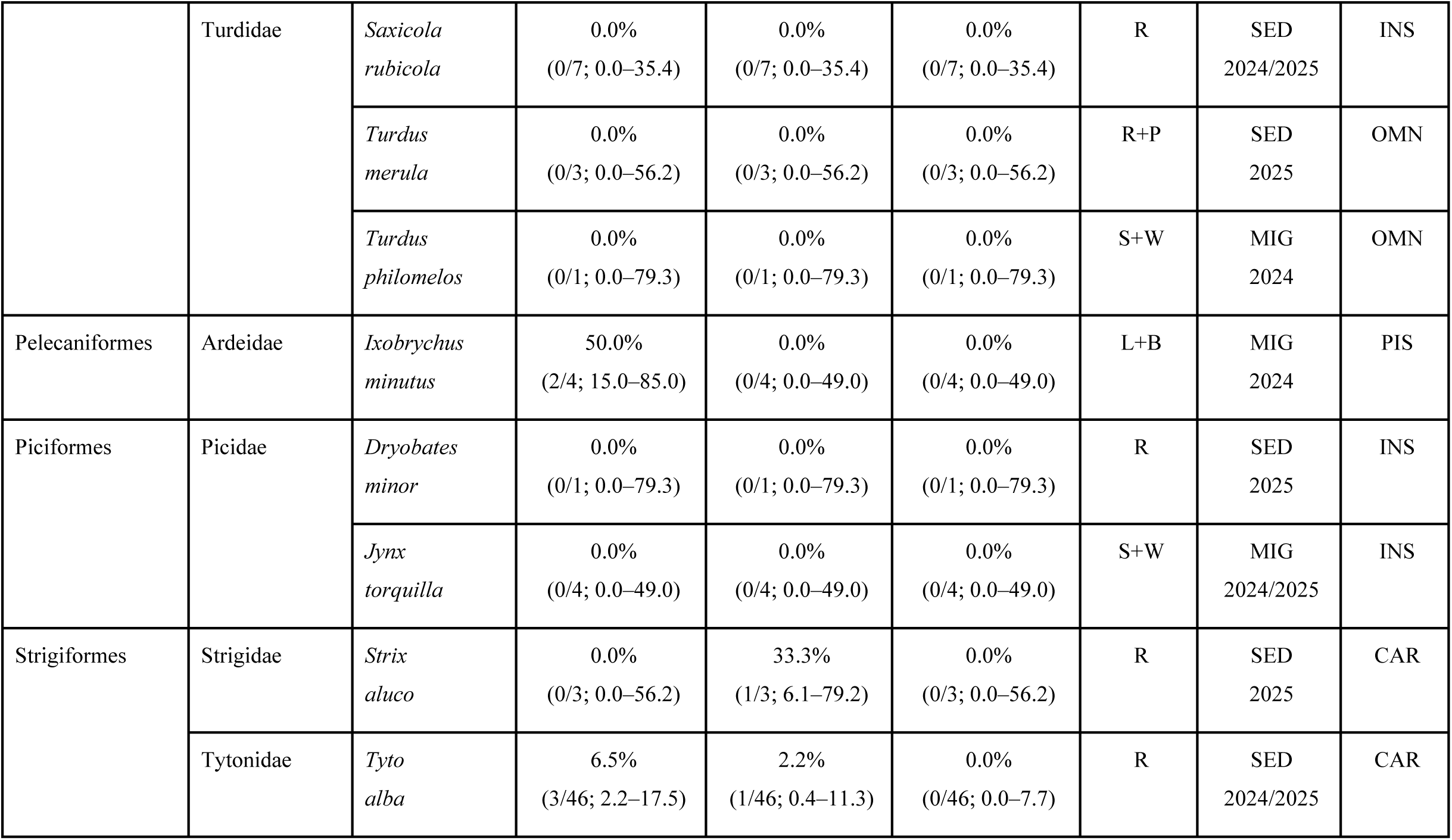
Summary of species traits and prevalence. The table presents species-level information including taxonomic identity, prevalence values, ecological traits, and associated classifications. For each species, prevalence represents the proportion of positive observations relative to the total number of individuals sampled, expressed as n/N with 95% confidence intervals (95% CI). USUV and WNV prevalences correspond to single infections only, whereas co-infection refers exclusively to simultaneous detection of both viruses in the same individual. Migration status is coded as follows: R = Resident; P = Partial migrant; S = Short-distance migrant; L = Long-distance migrant; W = Wintering; B = Breeding. SED/MIG 2024/2025 indicates the study-level classification of species as sedentary (SED) or migratory (MIG), and the year(s) of sampling (2024, 2025, or both). Diet category is coded as follows: INS = Insectivorous; INV = Invertivorous; OMN = Omnivorous; GRA = Granivorous; FRU = Frugivorous; CAR = Carnivorous; PIS = Piscivorous; m = migration; b = breeding; w = winter.

## Bibliography

1. Zannoli S, Sambri V. West Nile Virus and Usutu Virus Co-Circulation in Europe: Epidemiology and Implications. Microorganisms. 2019;7(7):184. doi:10.3390/microorganisms7070184

2. Clé M, Beck C, Salinas S, Lecollinet S, Gutierrez S, Van de Perre P, et al. Usutu virus: A new threat? Epidemiol Infect. 2019;147:e232. doi:10.1017/S0950268819001213

3. Simonin Y. Circulation of West Nile Virus and Usutu Virus in Europe: Overview and Challenges. Viruses. 2024;16(4):599. doi:10.3390/v16040599

4. Cadar D, Simonin Y. Human Usutu Virus Infections in Europe: A New Risk on Horizon? Viruses. 2022;15(1):77. doi:10.3390/v15010077

5. Sule WF, Oluwayelu DO, Hernández-Triana LM, Fooks AR, Venter M, Johnson N. Epidemiology and ecology of West Nile virus in sub-Saharan Africa. Parasit Vectors. 2018;11(1):414. doi:10.1186/s13071-018-2998-y

6. Engel D, Jöst H, Wink M, Börstler J, Bosch S, Garigliany MM, et al. Reconstruction of the evolutionary history and dispersal of Usutu virus, a neglected emerging arbovirus in Europe and Africa. mBio. 2016;7(1):1–12. doi:10.1128/mBio.01938-15

7. Tinto B, Kaboré DPA, Kagoné TS, Constant O, Barthelemy J, Kiba-Koumaré A, et al. Screening of Circulation of Usutu and West Nile Viruses: A One Health Approach in Humans, Domestic Animals and Mosquitoes in Burkina Faso, West Africa. Microorganisms. 2022;10(10):2016. doi:10.3390/microorganisms10102016

8. Berneck BS, Rockstroh A, Barzon L, Sinigaglia A, Vocale C, Landini MP, et al. Serological differentiation of West Nile virus- and Usutu virus-induced antibodies by envelope proteins with modified cross-reactive epitopes. Transbound Emerg Dis. 2022;69(5):2779–87. doi:10.1111/tbed.14429

9. Siljic M, Sehovic R, Jankovic M, Stamenkovic G, Loncar A, Todorovic M, et al. Evolutionary dynamics of Usutu virus: Worldwide dispersal patterns and transmission dynamics in Europe. Front Microbiol. 2023;14:1145981. doi:10.3389/fmicb.2023.1145981

10. Prat M, Jeanneau M, Rakotoarivony I, Duhayon M, Simonin Y, Savini G, et al. Virulence and transmission vary between Usutu virus lineages in Culex pipiens. PLoS Negl Trop Dis. 2024;18(6):e0012295. doi:10.1371/journal.pntd.0012295

11. Clé M, Constant O, Barthelemy J, Desmetz C, Martin MF, Lapeyre L, et al. Differential neurovirulence of Usutu virus lineages in mice and neuronal cells. J Neuroinflammation. 2021;18(1):11. doi:10.1186/s12974-020-02060-4

12. Koch RT, Erazo D, Folly AJ, Johnson N, Dellicour S, Grubaugh ND, et al. Genomic epidemiology of West Nile virus in Europe. One Health. 2023;18:100664. doi:10.1016/j.onehlt.2023.100664

13. Barzon L, Papa A, Lavezzo E, Franchin E, Pacenti M, Sinigaglia A, et al. Phylogenetic characterization of Central/Southern European lineage 2 West Nile virus: Analysis of human outbreaks in Italy and Greece, 2013-2014. Clin Microbiol Infect. 2015;21(12):1122.e1-10. doi:10.1016/j.cmi.2015.07.018

14. ECDC. Epidemiological update: West Nile virus transmission season in Europe, 2018 [Internet]. [cited 2026 Mar 25]. Available from: https://www.ecdc.europa.eu/en/news-events/epidemiological-update-west-nile-virus-transmission-season-europe-2018

15. ECDC. Historical data on local transmission in Europe for West Nile virus [Internet]. [cited 2026 Mar 25]. Available from: https://www.ecdc.europa.eu/en/west-nile-fever/surveillance-and-disease-data/historical

16. Bouchez-Zacria M, Calenge C, Villers A, Lecollinet S, Gonzalez G, Quintard B, et al. Relevance of the synergy of surveillance and populational networks in understanding the Usutu virus outbreak within common blackbirds (Turdus merula) in Metropolitan France, 2018. Peer Community Journal. 2025;5:e9. doi:10.24072/pcjournal.513

17. de Bruin E, van Irsel J, Chandler F, Kohl R, van de Voorde T, van der Linden A, et al. Usutu Virus Antibody Dynamics in Naturally Infected Blackbirds, the Netherlands, 2016-2018. Emerg Infect Dis. 2025;31(6):1244–6. doi:10.3201/eid3106.241744

18. Ziegler U, Lühken R, Keller M, Cadar D, Van Der Grinten E, Michel F, et al. West Nile virus epizootic in Germany, 2018. Antiviral Res. 2019;162:39–43. doi:10.1016/j.antiviral.2018.12.005

19. Michel F, Sieg M, Fischer D, Keller M, Eiden M, Reuschel M, et al. Evidence for West Nile Virus and Usutu Virus Infections in Wild and Resident Birds in Germany, 2017 and 2018. Viruses. 2019;11(7):674. doi:10.3390/v11070674

20. Münger E, Atama NC, van Irsel J, Blom R, Krol L, van Mastrigt T, et al. One Health approach uncovers emergence and dynamics of Usutu and West Nile viruses in the Netherlands. Nat Commun. 2025;16(1):7883. doi:10.1038/s41467-025-63122-w

21. Bakonyi T, Gould EA, Kolodziejek J, Weissenböck H, Nowotny N. Complete genome analysis and molecular characterization of Usutu virus that emerged in Austria in 2001: Comparison with the South African Strain SAAR-1776 and other flaviviruses. Virology. 2004;328(2):301–10. doi:10.1016/j.virol.2004.08.005

22. Chvala S, Bakonyi T, Bukovsky C, Meister T, Brugger K, Rubel F, et al. Monitoring of Usutu virus activity and spread by using dead bird surveillance in Austria, 2003-2005. Vet Microbiol. 2007;122(3–4):237–45. doi:10.1016/j.vetmic.2007.01.029

23. Rijks JM, Kik M, Slaterus R, Foppen R, Stroo A, Ijzer J, et al. Widespread Usutu virus outbreak in birds in The Netherlands, 2016. Euro Surveill. 2016;21(45):30391. doi:10.2807/1560-7917.ES.2016.21.45.30391

24. Folly AJ, Lawson B, Lean FZX, McCracken F, Spiro S, John SK, et al. Detection of Usutu virus infection in wild birds in the United Kingdom, 2020. Euro Surveill. 2020;25(41):2001732. doi:10.2807/1560-7917.ES.2020.25.41.2001732

25. Constant O, Bollore K, Clé M, Barthelemy J, Foulongne V, Chenet B, et al. Evidence of exposure to USUV and WNV in zoo animals in France. Pathogens. 2020;9(12):1005. doi:10.3390/pathogens9121005

26. Lauriano A, Rossi A, Galletti G, Casadei G, Santi A, Rubini S, et al. West Nile and Usutu Viruses’ Surveillance in Birds of the Province of Ferrara, Italy, from 2015 to 2019. Viruses. 2021;13(7):1367. doi:10.3390/v13071367

27. Tamba M, Bonilauri P, Galletti G, Casadei G, Santi A, Rossi A, et al. West Nile virus surveillance using sentinel birds: results of eleven years of testing in corvids in a region of northern Italy. Front Vet Sci. 2024;11:1407271. doi:10.3389/fvets.2024.1407271

28. Schopf F, Sadeghi B, Bergmann F, Fischer D, Rahner R, Müller K, et al. Circulation of West Nile virus and Usutu virus in birds in Germany, 2021 and 2022. Infect Dis. 2025;57(3):256–77. doi:10.1080/23744235.2024.2419859

29. Lecollinet S, Leblond A, Durand B, Zientara S, Ponçon N. Le virus West Nile : bilan de la situation en Europe et point sur la surveillance en France. Bulletin épidémiologique, santé animale et alimentation. 2012;(49):32–4.

30. Vilibic-Cavlek T, Savic V, Petrovic T, Toplak I, Barbic L, Petric D, et al. Emerging Trends in the Epidemiology of West Nile and Usutu Virus Infections in Southern Europe. Frontiers in veterinary science. 2019 Dec;6. doi:10.3389/FVETS.2019.00437

31. Csank T, Drzewnioková P, Korytár Ľ, Major P, Gyuranecz M, Pistl J, et al. A Serosurvey of Flavivirus Infection in Horses and Birds in Slovakia. Vector Borne Zoonotic Dis. 2018;18(4):206–13. doi:10.1089/vbz.2017.2216

32. Rizzoli A, Jiménez-Clavero MA, Barzon L, Cordioli P, Figuerola J, Koraka P, et al. The challenge of West Nile virus in Europe: knowledge gaps and research priorities. Euro Surveill. 2015;20(20):21135. doi:10.2807/1560-7917.ES2015.20.20.21135

33. Constant O, Gil P, Barthelemy J, Bolloré K, Foulongne V, Desmetz C, et al. One Health surveillance of West Nile and Usutu viruses: a repeated cross-sectional study exploring seroprevalence and endemicity in Southern France, 2016 to 2020. Euro Surveill. 2022;27(25):2200068. doi:10.2807/1560-7917.ES.2022.27.25.2200068

34. Simonin Y, Sillam O, Carles MJ, Gutierrez S, Gil P, Constant O, et al. Human Usutu virus infection with atypical neurologic presentation, Montpellier, France, 2016. Emerg Infect Dis. 2018;24(5):875–8. doi:10.3201/eid2405.171122

35. Eiden M, Gil P, Ziegler U, Rakotoarivony I, Marie A, Frances B, et al. Emergence of two Usutu virus lineages in Culex pipiens mosquitoes in the Camargue, France, 2015. Infect Genet Evol. 2018;61:151–4. doi:10.1016/j.meegid.2018.03.020

36. Vittecoq M, Lecollinet S, Jourdain E, Thomas F, Blanchon T, Arnal A, et al. Recent circulation of West Nile virus and potentially other closely related flaviviruses in Southern France. Vector Borne Zoonotic Dis. 2013;13(8):610–3. doi:10.1089/vbz.2012.1166

37. Beaubaton R, Revel J, Pigeyre L, Bollore K, Lepeule A, Mocq J, et al. Tracking the urban spread of Usutu virus in southern France: Detection across biological and environmental matrices. PLoS Negl Trop Dis. 2025;19(9):e0013506. doi:10.1371/journal.pntd.0013506

38. SPF. West nile virus [Internet]. [cited 2026 Mar 25]. Available from: https://www.santepubliquefrance.fr/maladies-et-traumatismes/maladies-a-transmission-vectorielle/west-nile-virus

39. Beck HE, Zimmermann NE, McVicar TR, Vergopolan N, Berg A, Wood EF. Present and future Köppen-Geiger climate classification maps at 1-km resolution. Sci Data. 2018;5:180214. doi:10.1038/sdata.2018.214

40. Pradier S, Sandoz A, Paul MC, Lefebvre G, Tran A, Maingault J, et al. Importance of wetlands management for West Nile Virus circulation risk, Camargue, Southern France. Int J Environ Res Public Health. 2014;11(8):7740–54. doi:10.3390/ijerph110807740

41. Knutie SA, Gotanda KM. A Non-invasive Method to Collect Fecal Samples from Wild Birds for Microbiome Studies. Microb Ecol. 2018;76(4):851–5. doi:10.1007/s00248-018-1182-4

42. Scaramozzino N, Crance JM, Jouan A, DeBriel DA, Stoll F, Garin D. Comparison of *Flavivirus* Universal Primer Pairs and Development of a Rapid, Highly Sensitive Heminested Reverse Transcription-PCR Assay for Detection of Flaviviruses Targeted to a Conserved Region of the NS5 Gene Sequences. J Clin Microbiol. 2001;39(5):1922–7. doi:10.1128/JCM.39.5.1922-1927.2001

43. Klitting R, Gondard M, Pezzi L, Migné CV, Bellone R, L’Ambert G, et al. Genomic Epidemiology of West Nile Virus in Paris. JAMA Netw Open. 2026;9(2):e2559588. doi:10.1001/jamanetworkopen.2025.59588

44. Mocq J, Raymond J, Bolloré K, Fossot A, Beaubaton R, Lepeule A, et al. Early-warning surveillance of West Nile and Usutu viruses in water and mosquito excreta using digital PCR. bioRxiv preprint. doi: 10.64898/2026.02.01.703130

45. Vilibic-Cavlek T, Petrovic T, Savic V, Barbic L, Tabain I, Stevanovic V, et al. Epidemiology of Usutu Virus: The European Scenario. Pathogens. 2020;9(9):699. doi:10.3390/pathogens9090699

46. Angeloni G, Bertola M, Lazzaro E, Morini M, Masi G, Sinigaglia A, et al. Epidemiology, surveillance and diagnosis of Usutu virus infection in the EU/EEA, 2012 to 2021. Euro Surveill. 2023;28(33):2200929. doi:10.2807/1560-7917.ES.2023.28.33.2200929

47. Tolsá MJ, García-Peña GE, Rico-Chávez O, Roche B, Suzán G. Macroecology of birds potentially susceptible to West Nile virus. Proc Biol Sci. 2018;285(1893):20182178. doi:10.1098/rspb.2018.2178

48. Rodríguez-Valencia V, Olive M, Le Goff G, Faisse M, Bourel M, L’Ambert G, et al. Host-feeding preferences of Culex pipiens and its potential significance for flavivirus transmission in the Camargue, France. Med Vet Entomol. 2025;39(3):614–25. doi:10.1111/mve.12802

49. Jourdain E, Zeller HG, Sabatier P, Murri S, Kayser Y, Greenland T, et al. Prevalence of West Nile Virus Neutralizing Antibodies in Wild Birds from the Camargue Area, Southern France. J Wildl Dis. 2008;44(3):766–71. doi:10.7589/0090-3558-44.3.766

50. Roiz D, Vázquez A, Ruiz S, Tenorio A, Soriguer R, Figuerola J. Evidence that Passerine Birds Act as Amplifying Hosts for Usutu Virus Circulation. EcoHealth. 2019;16(4):734– 42. doi:10.1007/s10393-019-01441-3

51. Nemeth N, Young G, Ndaluka C, Bielefeldt-Ohmann H, Komar N, Bowen R. Persistent West Nile virus infection in the house sparrow (Passer domesticus). Arch Virol. 2009;154(5):783–9. doi:10.1007/s00705-009-0369-x

52. Komar N, Panella NA, Burkhalter KL. Focal amplification and suppression of West Nile virus transmission associated with communal bird roosts in northern Colorado. J Vector Ecol. 2018;43(2):220–34. doi:10.1111/jvec.12306

53. Pérez-Ramírez E, Llorente F, Jiménez-Clavero M. Experimental infections of wild birds with West Nile Virus. Viruses. 2014;6(2):752–81. doi:10.3390/v6020752

54. SPF. Virus West Nile en France hexagonale. Bilan 2024. [Internet]. [cited 2026 Mar 25]. Available from: https://www.santepubliquefrance.fr/maladies-et-traumatismes/maladies-a-transmission-vectorielle/west-nile-virus/documents/bulletin-national/virus-west-nile-en-france-hexagonale.-bilan-2024

55. Bahuon C, Marcillaud-Pitel C, Bournez L, Leblond A, Beck C, Hars J, et al. West Nile virus epizootics in the Camargue (France) in 2015 and reinforcement of surveillance and control networks. Rev Sci Tech. 2016;35(3):811–24. doi:10.20506/rst.35.3.2571

56. SPF. Chikungunya, dengue, Zika et virus West-Nile en Occitanie. Bulletin du 20 novembre 2025. [Internet]. [cited 2026 Apr 10]. Available from: https://www.santepubliquefrance.fr/regions/occitanie/documents/bulletin-regional/2025/chikungunya-dengue-zika-et-virus-west-nile-en-occitanie.-bulletin-du-20-novembre-2025

57. RESPE. Appel à vigilance Fièvre de West Nile - 16 septembre 2025 [Internet]. 2025 [cited 2026 Apr 10]. Available from: https://respe.net/appel-a-vigilance-fievre-de-west-nile-16-septembre-2025/

58. ECDC. Monthly updates: Seasonal surveillance in humans and animals in 2025 for West Nile virus [Internet]. [cited 2026 Mar 25]. Available from: https://www.ecdc.europa.eu/en/infectious-disease-topics/west-nile-virus-infection/surveillance-and-disease-data/monthly-updates

59. ECDC. Weekly updates: Seasonal surveillance in humans in 2025 for West Nile virus [Internet]. [cited 2026 Mar 25]. Available from: https://www.ecdc.europa.eu/en/west-nile-fever/surveillance-and-disease-data/disease-data-ecdc

60. Calistri P, Giovannini A, Hubalek Z, Ionescu A, Monaco F, Savini G, et al. Epidemiology of West Nile in Europe and in the Mediterranean Basin. Open Virol J. 2010;4(2):29–37. doi:10.2174/1874357901004020029

61. Fros JJ, Miesen P, Vogels CB, Gaibani P, Sambri V, Martina BE, et al. Comparative Usutu and West Nile virus transmission potential by local Culex pipiens mosquitoes in north-western Europe. One Health. 2015;1:31–6. doi:10.1016/j.onehlt.2015.08.002

62. Folly AJ, Sewgobind S, Hernández-Triana LM, Mansfield KL, Lean FZX, Lawson B, et al. Evidence for overwintering and autochthonous transmission of Usutu virus to wild birds following its redetection in the United Kingdom. Transbound Emerg Dis. 2022;69(6):3684–92. doi:10.1111/tbed.14738

63. Schmidt V, Cramer K, Böttcher D, Heenemann K, Rückner A, Harzer M, et al. Usutu virus infection in aviary birds during the cold season. Avian Pathol. 2021;50(5):427–35. doi:10.1080/03079457.2021.1962003

64. Blanquer A, Rivas F, Gérardy M, Sarlet M, Moula N, Ziegler U, et al. Evaluation of Non-Vector Transmission of Usutu Virus in Domestic Canaries (Serinus canaria). Viruses. 2024;16(1):79. doi:10.3390/v16010079

65. Schilling M, Lawson B, Spiro S, Jagdev M, Vaux AGC, Bruce RC, et al. Vertical transmission in field-caught mosquitoes identifies a mechanism for the establishment of Usutu virus in a temperate country. Sci Rep. 2025;15(1):25252. doi:10.1038/s41598-025-09335-x

66. ECDC. Epidemiological update: West Nile virus transmission season in Europe, 2023 [Internet]. [cited 2026 Mar 25]. Available from: https://www.ecdc.europa.eu/en/news-events/epidemiological-update-west-nile-virus-transmission-season-europe-2023-0

67. ECDC, EFSA. Surveillance, prevention and control of West Nile virus and Usutu virus infections in the EU/EEA. 2023. doi:10.2903/sp.efsa.2023.EN-8242

68. Paz S. Climate change impacts on West Nile virus transmission in a global context. Phil Trans R Soc B. 2015;370(1665):20130561. doi:10.1098/rstb.2013.0561

69. Weidinger P, Kolodziejek J, Bakonyi T, Brunthaler R, Erdélyi K, Weissenböck H, et al. Different dynamics of Usutu virus infections in Austria and Hungary, 2017-2018. Transbound Emerg Dis. 2020;67(1):298–307. doi:10.1111/tbed.13351

70. Kilpatrick AM, Daszak P, Jones MJ, Marra PP, Kramer LD. Host heterogeneity dominates West Nile virus transmission. Proc R Soc B. 2006;273(1599):2327–33. doi:10.1098/rspb.2006.3575

71. Kilpatrick AM, Fonseca DM, Ebel GD, Reddy MR, Kramer LD. Spatial and Temporal Variation in Vector Competence of Culex pipiens and Cx. restuans Mosquitoes for West Nile Virus. Am J Trop Med Hyg. 2010;83(3):607–13. doi:10.4269/ajtmh.2010.10-0005

72. Wang H, Abbo SR, Visser TM, Westenberg M, Geertsema C, Fros JJ, et al. Competition between Usutu virus and West Nile virus during simultaneous and sequential infection of Culex pipiens mosquitoes. Emerg Microbes Infect. 2020;9(1):2642–52. doi:10.1080/22221751.2020.1854623

73. Hernández-Triana LM, Jeffries CL, Mansfield KL, Carnell G, Fooks AR, Johnson N. Emergence of West Nile virus lineage 2 in Europe: A review on the introduction and spread of a mosquito-borne disease. Front Public Health. 2014;2:271. doi:10.3389/fpubh.2014.00271

74. Beck C, Goffart IL, Franke F, Gonzalez G, Dumarest M, Lowenski S, et al. Contrasted Epidemiological Patterns of West Nile Virus Lineages 1 and 2 Infections in France from 2015 to 2019. Pathogens. 2020;9(11):908. doi:10.3390/pathogens9110908

75. Zecchin B, Fusaro A, Milani A, Schivo A, Ravagnan S, Ormelli S, et al. The central role of Italy in the spatial spread of USUTU virus in Europe. Virus Evolution. 2021;7(1):veab048. doi:10.1093/ve/veab048

76. Giglia G, Agliani G, Munnink BBO, Sikkema R, Mandara MT, Lepri E, et al. Pathology and Pathogenesis of Eurasian Blackbirds (Turdus merula) Naturally Infected with Usutu Virus. Viruses. 2021;13(8):1481. doi:10.3390/v13081481

77. Eidson M, Kramer L, Stone W, Hagiwara Y, Schmit K. Dead Bird Surveillance as an Early Warning System for West Nile Virus. Emerg Infect Dis. 2001;7(4):631–5. doi:10.3201/eid0704.010405

78. Mostashari F, Kulldorff M, Hartman JJ, Miller JR, Kulasekera V. Dead Bird Clusters as an Early Warning System for West Nile Virus Activity. Emerg Infect Dis. 2003;9(6):641– 6. doi:10.3201/eid0906.020794

